# Lasting alterations in monocyte and dendritic cell subsets in individuals after hospitalization for COVID-19

**DOI:** 10.1101/2022.07.15.500185

**Authors:** Francis R. Hopkins, Melissa Govender, Cecilia Svanberg, Johan Nordgren, Hjalmar Waller, Åsa Nilsdotter-Augustinsson, Anna J. Henningsson, Marie Hagbom, Johanna Sjöwall, Sofia Nyström, Marie Larsson

**Author notes:** Address correspondence to Marie Larsson, Molecular Medicine and Virology, Lab 1 Plan 13, Department of Biomedicine and clinical Sciences, Linköping University 58 185 Linköping, Sweden. These Authors contributed equally to this study (Francis R. Hopkins, Melissa Govender).

## Abstract

After more than two years the COVID-19 pandemic continues to burden healthcare systems and economies worldwide, and it is evident that long-term effects of the disease can persist for months post-recovery in some individuals. The activity of myeloid cells such as monocytes and dendritic cells (DC) is essential for correct mobilization of the innate and adaptive responses to a pathogen. Impaired levels and responses of monocytes and DC to SARS-CoV-2 is likely to be a driving force behind the immune dysregulation that characterizes severe COVID-19. Here, we followed, for 6-7 months, a cohort of COVID-19 patients hospitalized during the early waves of the pandemic. The levels and phenotypes of circulating monocyte and DC subsets were assessed to determine both the early and long-term effects of the SARS-CoV-2 infection. We found increased monocyte levels that persisted for 6-7 months, mostly attributed to elevated levels of classical monocytes. While most DC subsets recovered from an initial decrease, we found elevated levels of cDC2/cDC3 at the 6-7 month timepoint. Analysis of functional markers on monocytes and DC revealed sustained reduction in PD-L1 expression but increased CD86 expression across almost all cell types examined. Finally, viral load and CRP correlated to the appearance of circulating antibodies and levels of circulating DC and monocyte subsets, respectively. By elucidating some of the long-term effects that SARS-CoV-2 infection has on these key innate myeloid cells, we have shed more light on how the immune landscape remains affected in the months following severe COVID-19.

## INTRODUCTION

Since coronavirus disease 2019 (COVID-19) was declared a pandemic on March 11 2020, great progress has been made in combating the disease. To date, more than ten vaccines have been developed and granted emergency use listing by the World Health Organization ^1^. As of July 2022, the mortality from the disease stands at 6.3 million deaths, with 550 million cumulative cases worldwide ^2^, although with different standards of testing worldwide, the true human cost of the pandemic is likely to be much higher. The evolution of new variants indicate that the pandemic is still far from over.

The innate and adaptive immune responses during severe acute respiratory syndrome coronavirus 2 (SARS-CoV-2) infection have been extensively studied. Different pathways, markers, and factors of interest for drug targets or vaccine development have been identified and provide a better understanding of the impact of COVID-19 on the human host. While the various T cell responses in COVID-19 have been delineated ^3, 4^ there is still much unknown about other immune cells and their role in the disease.

The myeloid cell compartment consists of a variety of innate immune cell populations including dendritic cells (DC) and monocytes ^5^. Monocytes can be divided into different subsets based on their expression levels of CD14 and CD16 ^6^. Classical monocytes (cMo, CD14^+^CD16^-^) are highly phagocytic and can upregulate proteins associated with anti-bacterial activity, while non-classical monocytes (ncMo, CD14^-/low^CD16^+^) have a less inflammatory phenotype and are associated with wound healing ^7^. Intermediate monocytes (iMo, CD14^+^CD16^+^) are not completely understood but display a capacity for both antigen presentation and inflammatory responses ^8, 9^. In addition to monocytes, there are also myeloid derived suppressor cells (MDSC) that are strongly immunosuppressive, which increase in numbers in settings such as chronic infections and cancers ^10^.

The pathogenic role of monocytes in respiratory viral infection has been demonstrated in mouse models of influenza A infection, where inflammatory monocytes drive the virus-induced damage to the lung ^11^. Alterations to the monocyte compartment occur during SARS-CoV-2 infection, with most findings showing strong inflammatory monocyte responses in severe disease ^12, 13^, whereas the opposite is found for mild disease ^14^. Indeed, it has been shown that classical monocytes are drivers of the cytokine storm that frequently proves fatal in SARS-CoV-2 infection ^15^.

The DC serve as a bridge between the innate and adaptive responses and are essential for the development of an adaptive response ^16^. Like monocytes, DC can be classified into different subsets with the ability to sense pathogens, produce cytokines and activate T cell responses ^17^. Plasmacytoid DC (pDC) are defined by surface expression of CD303 (BDCA-2) and are the main type I interferon (IFN) producers ^18, 19^. Conventional DC (cDC) are further divided into different subsets, with cDC1 having a high capacity to cross-present antigens and activate T helper (Th)1 responses ^17, 20^, while cDC2 activate a wider range of Th responses ^17^. Recent work has divided cDC2 further, based on their CD5 expression, into a CD5^high^ population mainly promoting Th1 responses, and a CD5^low^ population promoting Th2, Th17, Th22, and T regulatory cell responses ^21^. In addition, within the traditionally defined cDC2 population the most newly characterized populations are the two cDC3 populations ^17^.

Functionality of monocyte and DC subsets can be defined by their expression of surface markers such as HLA-DR, CD86, CCR7 and PD-L1. In COVID-19 patients there is evidence of lower HLA-DR expression on DC subtypes, but not monocytes, indicating an impaired ability to activate T cells. Increased PD-L1 expression is also indicative of an impairment of the effector function of DC ^22^.

Studies of mild and severe COVID-19 patients have demonstrated an early overall decrease in circulating DC populations, particularly the pDC ^23, 24^. Furthermore, in severe disease the different DC subsets are impaired in their ability to sense pathogens, present antigens, and stimulate T cell responses ^25-27^. The importance of a functional DC compartment is illustrated by the correlation between the activation of early and effective T cell responses, and a positive clinical outcome in SARS-CoV-2 infection ^28, 29^. So far, there have been few longitudinal studies investigating the effects of COVID-19 on myeloid cell subsets during the disease and after recovery ^30^, and more data is needed to provide better insight into the long-term effects of COVID-19.

Here, we have characterized the effects that severe COVID-19 exerts on circulating DC, monocytes, and MDSC, in patients that needed hospitalization, using spectral flow cytometry. Furthermore, the effects of several clinical markers on levels of myeloid cell subsets were assessed.

In accordance with previous studies, all circulating DC subsets were decreased at inclusion, during acute infection. While pDC and cDC1 subsequently returned to levels comparable to healthy controls, the cDC2 and cDC3 combined subset was significantly elevated at the 6-7 month time point. We further found the frequencies of different monocyte subsets to be altered. An initial increase of cMo and decrease of iMo and ncMo was observed, and this profile was maintained long-term for cMo and iMo in comparison to healthy controls. Additionally, the MDSC compartment remained elevated, confirming a long-term immunosuppressive environment post-COVID-19. In addition, there was an alteration of immunomodulatory markers, such as HLA-DR, CD86 and PD-L1, in all cell subsets. Together, our findings highlight sustained long-term alterations in monocyte and DC subsets after COVID-19, that could be linked to initial elevated circulating C-reactive protein (CRP) levels.

## MATERIALS AND METHODS

### Clinical data and study design

This study was approved by the Swedish Ethical Review Authority (Ethics No. 2020-02580). The hospitalized COVID-19 patients were included at the Clinic of Infectious Diseases and the Intensive Care Unit (ICU) at the Vrinnevi Hospital, Norrköping, Sweden. Furthermore, healthy COVID-19 negative controls were enrolled among the staff at the same hospital. For this study we used longitudinal samples from hospitalized COVID-19 patients (N=21; age range 32-83 years) and samples from healthy controls (N=16; age range 23-80 years). Sample collection from COVID-19 patients was performed at four timepoints throughout the study: at enrolment when the COVID-19 patients were admitted to the Department of Infectious Diseases and after 2 weeks, 6 weeks, and 6-7 months post-enrolment. Both male and female participants were included, and all individuals had provided written informed consent prior to enrolment. This study was carried out in accordance with the International Conference on Harmonization Guidelines and the Declaration of Helsinki. Clinical data is described in **Table 1**.

**Table 1.** Clinical and-demographical characteristics of hospitalized COVID-19 patients.

| Variable | Clinical data | Reference range |
| --- | --- | --- |
| Number of patients | 21 |  |
| Age, median (range) | 56 (32–83) |  |
| Body mass index, median (range) | 30.7 (23.7–45.2) |  |
| Biological sex, % (N) | 38.1 F/61.9 M (8F/13M) |  |
| Days in hospital, median (range) | 9 (4–34) |  |
| ICU/pandemic Ward %, (N) | 14.3/85.7 (3/18) |  |
| Days with symptoms before inclusion, median (range) | 11 (5–30) |  |
| Spike IgG antibody positive at inclusion, % (N) | 76.2 (16) |  |
| Nucleocapsid IgG antibody positive at inclusion, % (N) | 76.2 (16) |  |
| Viral load at inclusion, median (range)(copies/ml) | 3589 (1071–111 x 10<sup>6</sup>) |  |
| Antiviral treatment, % (N) | 42.9 (9) |  |
| Corticosteroid treatment, % (N) | 76.2 (16) |  |
| No/oxygen/HFNOT:CPAP<sup>1</sup>/mechanical ventilation, % (N) | 4.8/38.1/52.3/4.8 (1/8/11/1) |  |
| Cardiovascular disease, % (N) | 52.4 (11) |  |
| Pulmonary disease, % (N) | 28.6 (6) |  |
| Diabetes mellitus, % (N) | 19 (4) |  |
| Two of the underlying conditions, % (N) | 19 (4) |  |
| Three of the underlying conditions, % (N) | 9.5 (2) |  |
| Disease score: moderate/severe, % (N) | 85.7 (18)/14.3 (3) |  |
| Smoker/snus, % (N) | 19 (4) |  |
| Previous history of smoking/snus, % (N) | 47.6 (10) |  |
| Leukocytes (x 10<sup>9</sup>/L), median (range) | 6.6 (3.5–20.4) | 3.5–8.8 |
| Thrombocytes (x 10<sup>9</sup>/L), median (range) | 275 (151–476) | 150–400 |
| Lymphocytes (x 10<sup>9</sup>/L), median (range) | 0.9 (0.2–2.8) | 1.1–4.8 |
| Monocytes (x 10<sup>9</sup>/L), median (range) | 0.4 (0.1–1.33) | 0.1–1 |
| Lactate dehydrogenase (μKat/L), median (range) | 7.4 (3.9–16) | >70 years <3.5, <70 years < 4.3 |
| C-reactive protein (mg/L), median (range) | 86 (14–295) | 0–10 |

<sup>1</sup>High flow nasal oxygen therapy (HFNOT), continuous positive airway pressure therapy (CPAP).
Diabetes mellitus, cardiovascular, and chronic pulmonary diseases defined by the individuals being medicated for these conditions.

Data regarding the clinical symptoms related to COVID-19 and general health status were collected from all participants included in the study. The information is summarized in **Table 1**. The severity of the disease was determined in the individuals hospitalized for COVID-19 as per the criteria defined by the National Institutes of Health ^31^, i.e., estimated with regards to maximum oxygen needed, and highest level of care provided. COVID-19 severity was classified as: mild (admitted to pandemic department, no oxygen supplementation), moderate (admitted to pandemic department, oxygen supplementation <5L/min), severe (admitted to pandemic department or intermediate care unit, oxygen need ≥5L/min, supplemented by high flow nasal oxygen therapy (HFNOT) and continuous positive airway pressure therapy (CPAP)) and critical (intensive care unit, with or without mechanical ventilation).

### Sample collection and processing

Blood samples were collected from study subjects in EDTA-treated Vacuette® tubes (Griener Bio-one GmbH, Kremsmünster, Austria) and peripheral blood mononuclear cells (PBMCs) were isolated by density gradient centrifugation (Ficoll-Paque, GE Healthcare, ThermoFisher). The PBMCs were washed and cryopreserved in freezing medium (8% DMSO in FBS) at -150°C until use. Sera were isolated from whole blood samples that had been collected in 3.5 ml Vacuette® tubes (Griener Bio-one GmbH, Kremsmünster, Austria), by centrifugation at 1000 g for 10 minutes at room temperature. Sera were frozen and stored at -80°C until testing.

### Spectral Flow cytometry

Cryopreserved PBMCs were thawed and washed in RPMI before resuspension in buffer (0.2% FBS in PBS) at concentration of 1 × 10^6^ cells/ml in FACS tubes (Falcon® Brand, VWR). Before addition of the cocktail of 21 monoclonal antibodies, cells were blocked with a cocktail of FcγR (1/15 dilution, Miltenyi), Novablock (Phitonex), and stained with live/dead violet viability dye (**Supplementary Table 1 for the antibodies and spectral flow reagents**), for 20 mins at 4°C. After washing, 30 μl of antibody cocktail was added to the cells, and the mix incubated for 30 minutes at 4°C. The stained PBMCs were washed and resuspended in 200 μl buffer and samples acquired using a Cytek Aurora (USA) spectral flow cytometer. Data was processed using OMIQ (California, USA).

### SARS-CoV-2 neutralizing antibodies

To measure neutralizing antibodies, against the SARS-CoV-2, we utilized the TECO® SARS-CoV-2 Neutralization Antibody Assay (TECOmedical AG, Sissach, Switzerland). The assay was performed according to the protocol provided by the manufacturer. Briefly, serum samples were diluted in sample buffer (1:10, 1:30, and 1:90) and incubated at 37°C for 30 minutes in a 96-well plate coated with ACE2 (provided in the kit). The plate was washed 3 times with diluted wash buffer (provided in the kit) and incubated with (S)-RBD–horseradish peroxidase conjugate for 15 minutes at 37°C. Finally, stop solution (provided in the kit) was added and optical density measured within 5 minutes at 450 nm (SpectraMax iD3 Molecular Devices, USA). The inhibition rate was calculated, and a positive value with the cutoff set at ≥20%.

### Data analysis and statistics

Data analysis and statistical calculations were performed with GraphPad Prism v9 (GraphPad Software, CA, USA). Differences among the study groups were analyzed with either unpaired, parametric T test with Welch’s correction, or Brown-Forsythe and Welch ANOVA tests, with no correction for multiple comparisons. In addition, bivariate analysis with Spearman’s correlation coefficient was performed. All differences with p values of <0.05 were considered statistically significant.

## RESULTS

### Study outline and clinical parameters

Patients suffering from COVID-19 and admitted to a hospital in Norrköping, Sweden were recruited into our study from May 2020 and throughout 2021, during and after the first and second waves of the pandemic in Sweden. Healthy controls who were confirmed SARS-CoV-2 negative by PCR and antibody tests, and had no prior history of COVID-19, were also recruited from within the hospital staff. Detailed clinical data were recorded for all individuals within the hospitalized cohort (**Table 1**). Whole blood, serum, and nasal swabs were collected at inclusion, and patients were followed up for further sample collection at 2 weeks, 6 weeks, and 6-7 months post-inclusion (**Figure 1A**). For this study, we have utilized longitudinal samples from 21 hospitalized COVID-19 patients and 16 healthy controls. Disease severity was defined according to the NIH guidelines, which are based largely on supplemental oxygen requirements ^31^. An initial assessment, at study inclusion, was made of an array of clinical parameters (**Figure 1B-C**), showing a significant decrease in lymphocytes, basophils, eosinophils, and neutrophils in the COVID-19 patients. The CRP and lactate dehydrogenase (LDH) levels were both significantly elevated in patients compared to healthy controls. All patients developed neutralizing SARS-CoV-2-specific antibodies that, in most individuals, lasted until the end of the study, even if the levels of neutralizing antibodies were significantly decreased at the 6-7 months timepoint compared to 2 and 6 weeks (**Figure 1D**). These findings are in agreement with previous publications ^32-36^. We performed correlation analysis of clinical parameters and found that CRP correlated with viral load at inclusion (**Supplementary Figure 1A**). Additional correlation analysis of clinical parameters and antibody levels revealed that anti-spike IgG levels and neutralizing antibody titers were negatively correlated with viral load at inclusion (**Supplementary Figure 1B**). The connection between viral load and both spike and nucleocapsid antibody levels has been shown previously ^37, 38^. Furthermore, there was negative correlation of neutralizing antibody titers and anti-spike IgG levels to CRP at inclusion (**Supplementary Figure 1C**). Following spectral flow data acquisition, lineage exclusion was performed to remove T cells, B cells, NK cells and basophils (**Figure 1E**), and remaining cells were defined as myeloid and used for further analysis of monocyte and DC populations. Major immune cell types from within PBMCs, and their changing distributions over time were visualized following dimensionality reduction analysis (**Figure 1F**).

**Figure 1.**
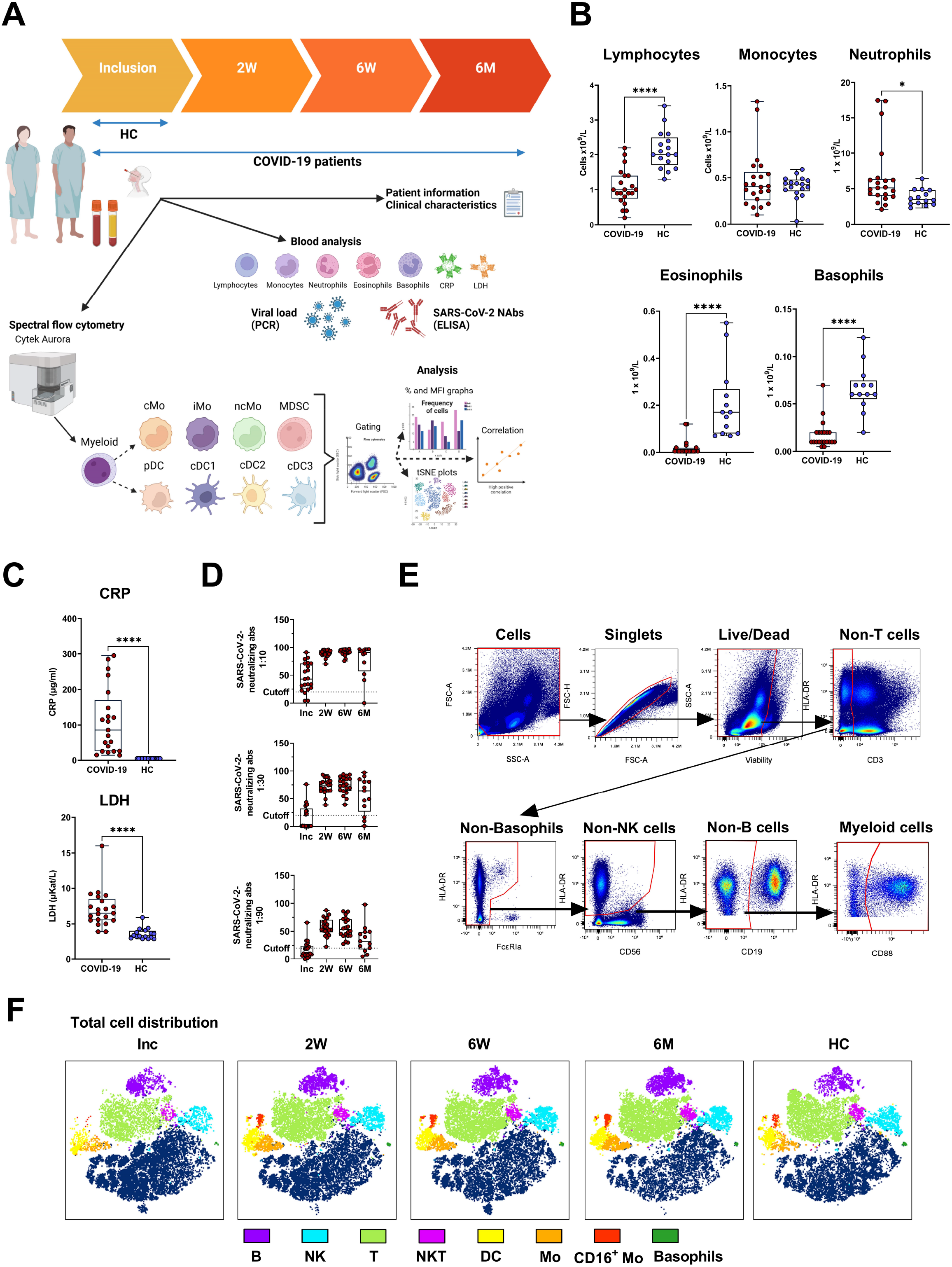
Study outline and experimental design. PBMCs and serum were collected from 21 COVID-19 patients and 16 healthy donors over a 6-7 month period. (**A**) Design of COVID cohort study illustrating timepoints for sample collection and experimental outputs, created with BioRender.com. Initial clinical assessments of (**B**) blood cell counts, (**C**) plasma C-reactive protein (CRP) and lactate dehydrogenase (LDH), and (**D**) neutralizing SARS-CoV-2 antibody titers. (**E**) Representative gating strategy to demonstrate live/dead gating and lineage exclusion, to leave myeloid cell populations for further analysis. (**F**) tSNE plots showing distinct clustering of all live cells, with 17 patients overlaid, and 16 controls overlaid. Data is represented as mean with 95% Cl, with significance of *p ≤ 0.05, ****p ≤ 0.0001, determined using Brown-Forsythe and Welch ANOVA tests. Inc = Study inclusion, 2W = 2 weeks, 6W = 6 weeks, 6M = 6-7 months, HC = healthy control. Mo = monocytes; cMo = classical iMo = intermediate, ncMo = non-classical, MDSC = myeloid-derived suppressor cells, DC = dendritic cells; pDC = plasmacytoid, cDC1/2/3 = conventional.

### Increased levels of monocytes in the mononuclear myeloid compartment

The proportion of total HLA-DR^+^ myeloid cells did not change significantly over the 6-7 months of the study (**Figure 2A**). Within the myeloid compartment in COVID-19 patients, monocytes were defined as CD88^+^ cells, with CD88 being exclusively expressed on blood monocytes ^39^. Monocytes increased significantly during the first 2 weeks after inclusion and reached a plateau that was significantly higher at 6-7 months post COVID-19 compared to controls (**Figure 2B**). The proportion of CD14^+^HLA-DR^-^ MDSC was raised compared to healthy controls at inclusion and at 2 weeks and 6-7 months follow up, but surprisingly, there was a significant dip at the 6 week time point (**Figure 2C**). Compared to the healthy controls there was a shift towards the non-monocytes, i.e., CD88^-^ cells, in the myeloid compartment following COVID-19 (**Figure 2D**). Together our results show that there is a sustained shift in the balance of cell types within the myeloid compartment (**Figure 2D**), despite the total level of myeloid cells remaining the same (**Figure 2A**).

**Figure 2.**
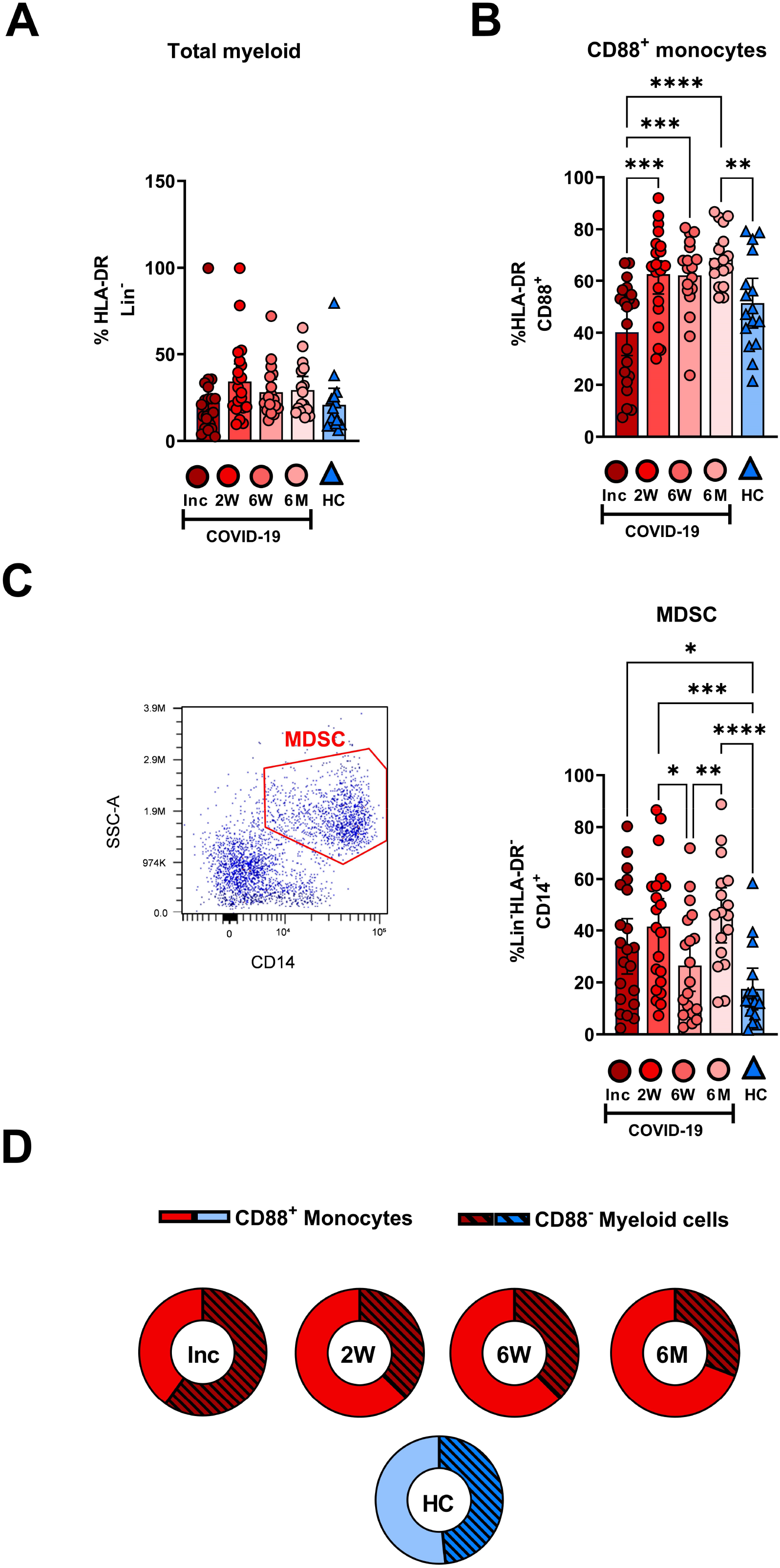
Dysregulation in the frequency of myeloid compartments persists for 6 months in hospitalized COVID-19 patients. PBMCs obtained from 21 COVID-19 patients and 16 healthy donors over a 6-7 month period were stained with monoclonal antibodies and assessed by flow cytometry to identify the myeloid cell subsets. Percentage of the cells comprising (**A**) the total HLA-DR^+^Lin^-^ myeloid compartment and (**B**) CD88^+^ monocytes. (**C**) Representative gating and percentage of CD14^+^HLA^-^DR-Lin^-^ MDSC. (**D**) Ratio of CD88^+^ monocytes to CD88^-^ myeloid cells. Data is represented as mean with 95% Cl, with significance of *p ≤ 0.05, **p ≤ 0.01, ***p ≤ 0.001, ****p ≤ 0.0001, determined using Brown-Forsythe and Welch ANOVA tests. Inc = Study inclusion, 2W = 2 weeks, 6W = 6 weeks, 6M = 6-7 months, HC = healthy control.

### Long term changes in classical and intermediate monocytes post COVID-19

The CD14^+^CD16^-^ cMo, CD14^+^CD16^+^ iMo and CD14^-/lo^CD16^+^ ncMo were examined to compare COVID-19 patients to healthy controls (**Figure 3**). The CD14^+^ and CD16^+^ cell levels within the myeloid compartment were initially low and increased over the 6-7 months (**Figure 3A**). The proportion of cMo at inclusion was significantly higher than in healthy controls and remained significantly elevated even after 6-7 months (**Figure 3B**). The iMo were reduced for the entire duration of the study (**Figure 3C**). A drastic reduction in ncMo at inclusion was observed in patients, but had already returned by 2 weeks to a level comparable to controls (**Figure 3C**). Together these data revealed that there was a sustained effect on the proportion of cMo and iMo in COVID-19 patients, while the change in ncMo was restored within 2 weeks.

**Figure 3.**
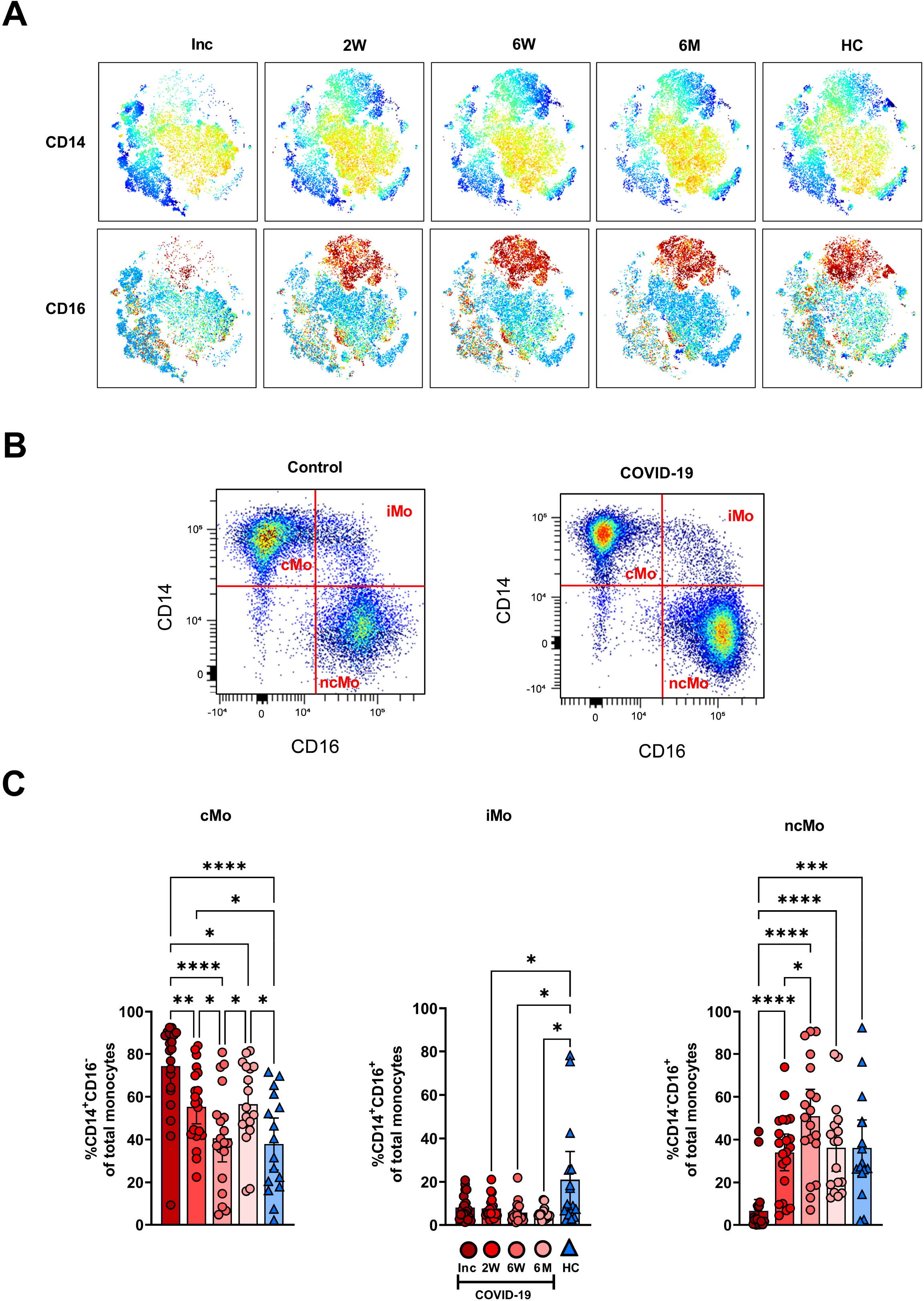
Classical and intermediate monocyte cells subsets remain altered in COVID-19 patients at 6 months. PBMCs were collected from COVID-19 patients that required hospitalization (N=21) and healthy controls (N=16) over a 6-7-month period. Cells were stained with antibodies for flow cytometry to define the monocyte subsets. (**A**) tSNE plots displaying myeloid cell clusters, colored according to their intensity of CD14 and CD16 expression. (**B**) Representative plots illustrating how monocyte subsets were defined. (**C**) Frequencies of the CD14^+^CD16^-^ cMo, CD14^+^CD16^+^ iMo, and CD14^-^CD16^-^ ncMo. Data is represented as mean with 95% Cl, with significance of *p ≤ 0.05, **p ≤ 0.01, ***p ≤ 0.001, ****p ≤ 0.0001, determined using Brown-Forsythe and Welch ANOVA tests. Inc = Study inclusion, 2W = 2 weeks, 6W = 6 weeks, 6M = 6-7 months, HC = healthy control.

### Circulating conventional DC subsets and plasmacytoid DC levels recovered after initial depletion

We assessed the different circulating DC subsets (**Figure 4A-B**) and found decreased levels of all DC subsets in the patients at study inclusion. The proportion of pDC were significantly reduced in COVID-19 patients at inclusion and that these cells started to recover already at two weeks (**Figure 4A**). The pDC fraction amongst the CD88^-^ myeloid cell compartment clearly increased until it reached a similar level to healthy controls (**Figure 4C**). The level had increased significantly by 2 weeks but did not fully return to normal until 6 weeks (**Figures 4A-C-D**). Next, we explored the different cDC subsets in COVID-19 patients. Among the cDC, CD141^+^ cDC1 constitute a very small fraction ^17^. This DC subset (**Figure 4A**) was significantly reduced at inclusion but throughout the 6-7 months studied, returned to a level comparable to controls (**Figure 5A**). Here, we analyzed cDC2 and cDC3 as one combined population due to the contrasting methods used by different groups to further define distinct cDC subpopulations within classical cDC2. While the proportion of the cumulative cDC2 and cDC3 subset was significantly reduced at inclusion, this recovered over the study period and even significantly surpassed the levels found in healthy controls and at 6-7 months (**Figure 5B**). Following this, cDC2 and cDC3 were further divided based on their expression of CD5 and CD14, respectively. Interestingly, neither the CD5^+^ cDC2 nor the CD5^*-*^ cDC2 were significantly altered at any of the study timepoints (**Figure 5C**). There was a significant increase in CD14^+^ cDC3 at the 2 week and 6-7 month time points, but no change in the CD14^-^ cDC3 (**Figure 5D**). Together, we found that both pDC and cDC1 recovered from initial reductions during the study. Additionally, there were major alterations over time in the combined cDC2 and cDC3 population, though the proportions of the subpopulations within these DC subsets were unaffected.

**Figure 4.**
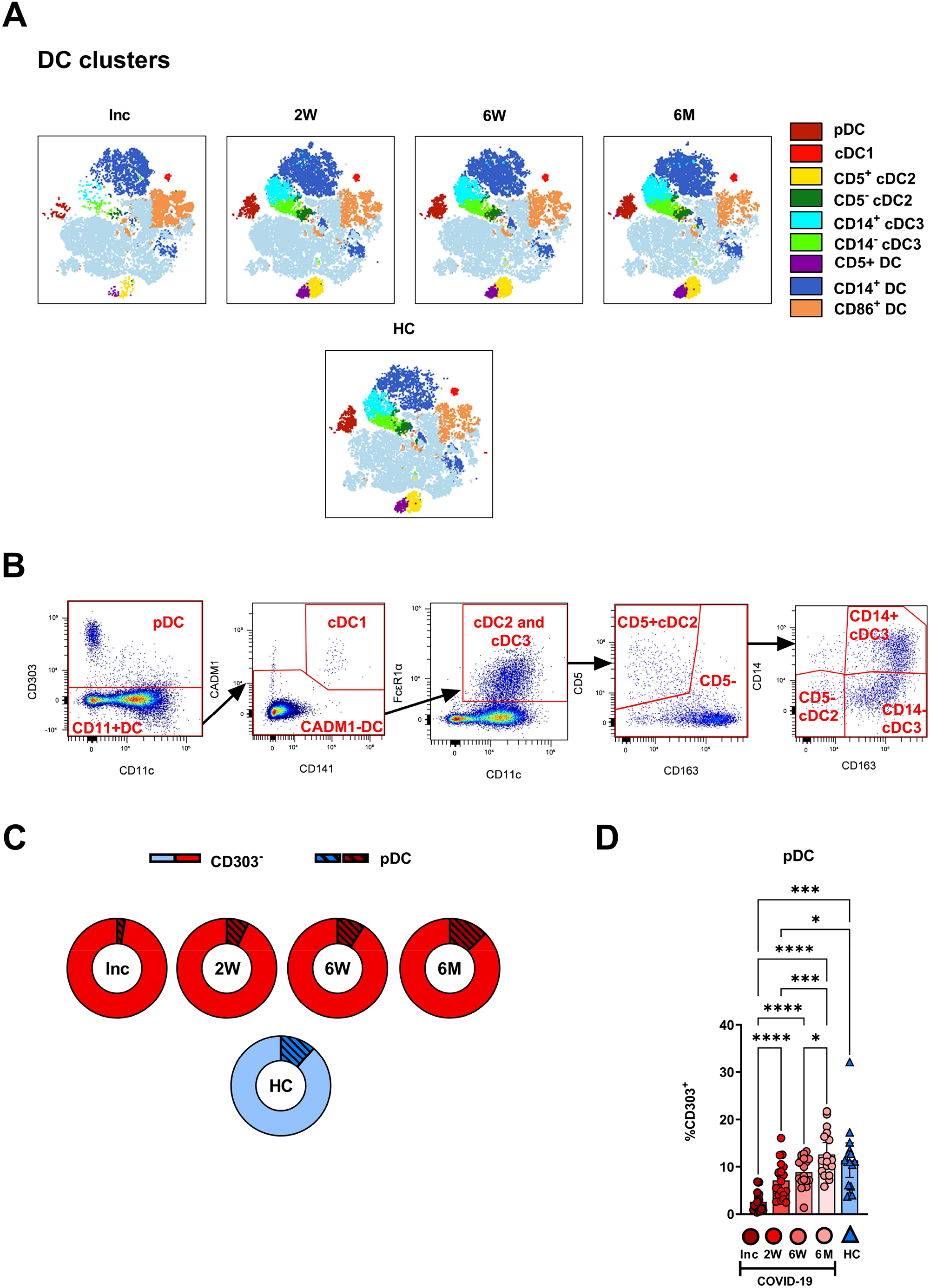
Plasmacytoid DCs in COVID-19 patients are severely depleted in blood during COVID-19. PBMCs collected from hospitalized COVID-19 patients (N = 21) and healthy controls (N = 16), were stained for flow cytometry to observe the effect of SARS-CoV-2 on the overall DC compartment. (**A**) tSNE plots illustrating the changing distribution of DC within CD88^-^ myeloid cells. (**B**) Gating strategy for DC subsets. (**C**) Ratio of CD303^-^ cells to pDC during COVID-19. (**D**) Percentages CD303^+^ pDC. Data are presented as mean with 95% Cl, with significance of *p ≤ 0.05, ***p ≤ 0.001, **** p ≤ 0.0001, determined using Brown-Forsythe and Welch ANOVA tests. Inc = Study inclusion, 2W = 2 weeks, 6W = 6 weeks, 6M = 6-7 months, HC = healthy control.

**Figure 5.**
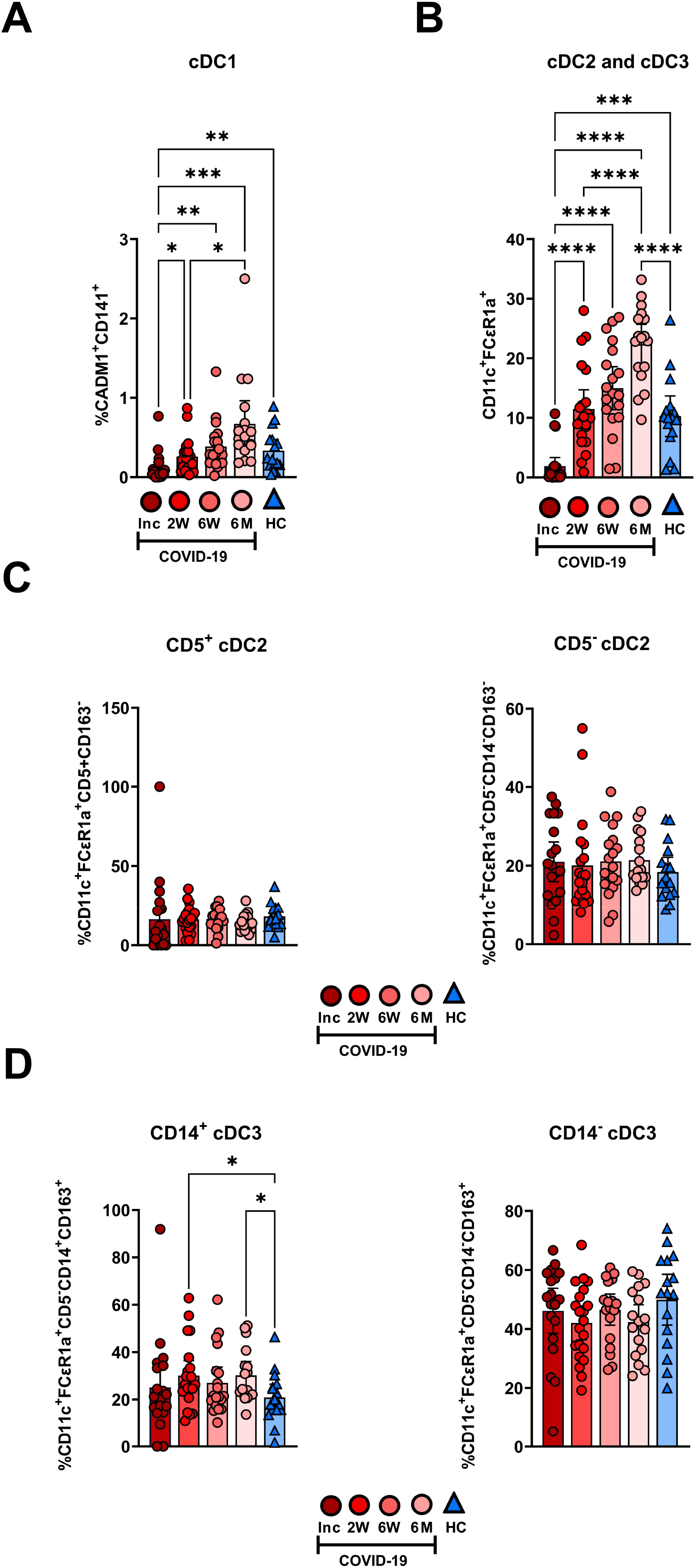
Initial depletion of classical DC populations during COVID-19. PBMCs from hospitalized COVID-19 patients (N = 21) and healthy donors (N = 16), were stained for flow cytometry to observe changes in the composition of the cDC compartment during COVID-19. Proportions of (**A**) CADM1^+^CD141^+^ cDC1, (**B**) CD11c^+^FCεR1a^+^ cDC2 and cDC3 combined, (**C**) CD5^+^CD163^-^ cDC2 and CD5^-^CD163^-^ cDC2, and (**D**) CD5^-^CD14^+^CD163^+^ cDC3 and CD5^-^CD14^+^CD163^+^ cDC3 are shown. Data is presented as mean with 95% Cl, with significance of *p ≤ 0.05, **p ≤ 0.01, ***p ≤ 0.001, ****p ≤ 0.0001, determined using Brown-Forsythe and Welch ANOVA tests. Inc = Study inclusion, 2W = 2 weeks, 6W = 6 weeks, 6M = 6-7 months, HC = healthy control.

### Prolonged decrease in PD-L1 and elevated CD86 and HLA-DR expression levels on monocyte subsets

The expression levels of functional surface markers were examined on monocyte subsets and MDSC (**Figure 6A, Supplementary Figure 2**). The clearest pattern was the decreased PD-L1 expression, which was observed at all time points on the monocyte subsets and the MDSC (**Figure 6A-B, Supplementary Figure 2**). In addition, there were increased CD86 and HLA-DR levels from the 2 week timepoint onwards on the monocyte subsets (**Figure 6A-C-D**). The iMo were selected to represent the PD-L1 expression pattern seen across all monocytic cell types, which was significantly reduced at all time points compared to healthy controls (**Figure 6B**). A detailed look at CD86 expression levels revealed a significant increased expression from 2 weeks onwards in iMo and ncMo, from 6 weeks onwards in MDSC, but not until 6 months in cMo (**Figure 6C**). HLA-DR expression was initially reduced in cMo before reverting to levels found in healthy controls at the 2 week time point. At the same time, there were increased HLA-DR levels from the 2 week time point onwards in iMo. HLA-DR expression on ncMo was similar at all time points (**Figure 6D**).

**Figure 6.**
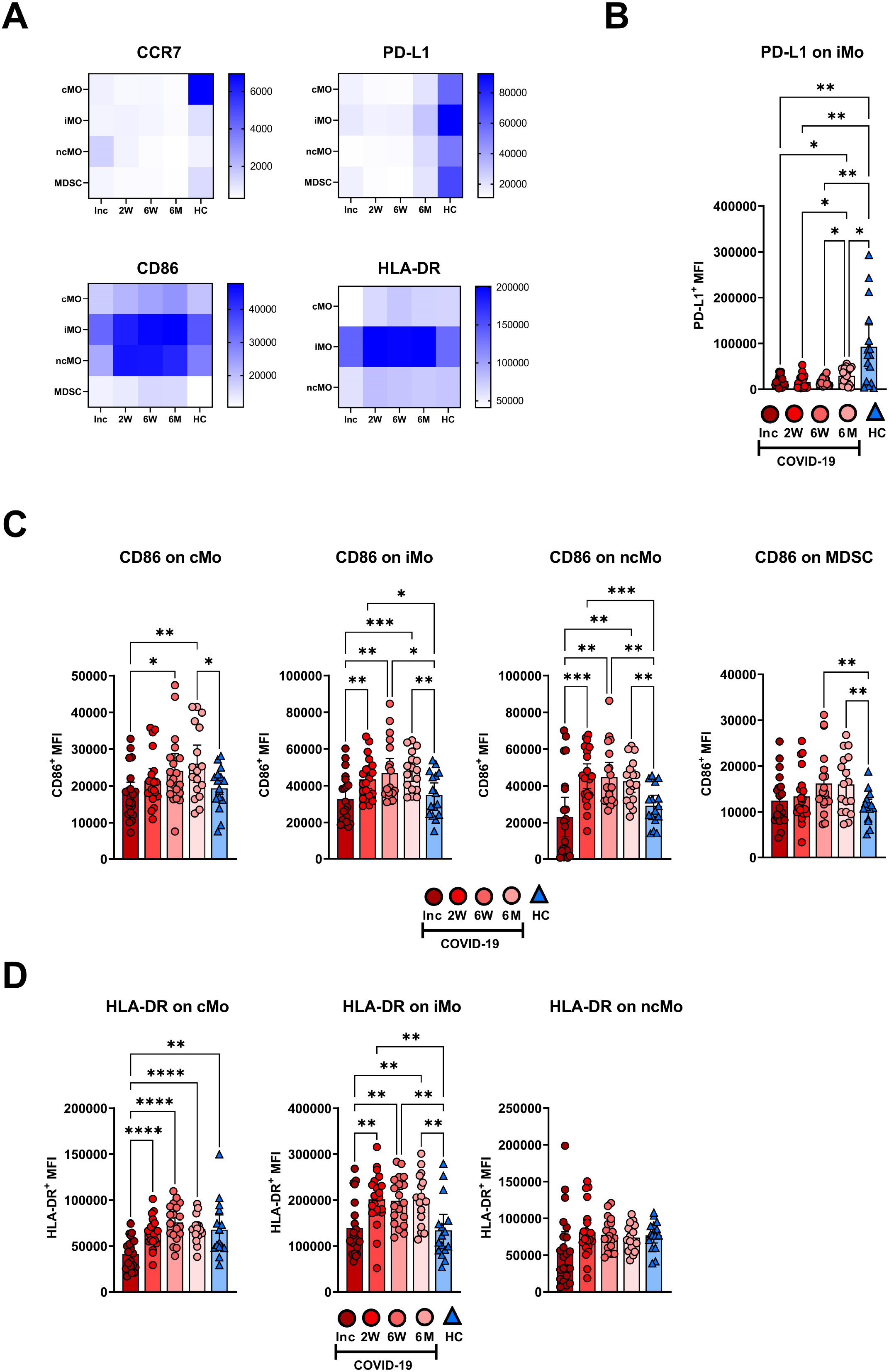
Long-term changes in the phenotype of blood monocyte subsets and myeloid derived suppressor cells in COVID-19 patients. PBMCs were collected from hospitalized COVID-19 patients (N=21) and healthy controls (N = 16) over a 6-7 month period. (**A**) Heatmaps showing the mean fluorescence intensity (MFI) of CCR7, PD-L1, CD86, and HLA-DR on monocyte subsets and MDSC, from all donors. (**B**) Representative graph of PD-L1 on iMo. MFI of (**C**) CD86 and (**D**) HLA-DR on monocyte subsets and MDSC. Data is represented as mean with 95% Cl, with significance of *p ≤ 0.05, **p ≤ 0.01, ***p ≤ 0.001, ****p ≤ 0.0001., determined using Brown-Forsythe and Welch ANOVA tests. Inc = Study inclusion, 2W = 2 weeks, 6W = 6 weeks, 6M = 6-7 months, HC = healthy control.

### Prolonged decrease in PD-L1 expression on all DC subsets and elevated CD86 levels on conventional DC subsets

The DC subsets were assessed for the expression levels of functional surface markers (**Figure 7A-B, Supplementary Figure 3**). There was a clear reduction in CD83 at 6 months and a decrease in PD-L1 at all time points, across all DC subsets (**Figure 7A**). The pDC and cDC3 were found to have decreased expression of CCR7 at the 6-7 month time point (**Figure 7A, Supplementary Figure 3**). PD-L1 was significantly decreased during the entire study across all DC subtypes studied (**Figure 7A, Supplementary Figure 3**). CD86 expression on pDC was significantly increased at inclusion but returned, by 2 weeks, to a comparable level to controls and remained so for the remainder of the 6-7 months post-COVID-19. On cDC1, CD86 was increased at both inclusion and the 6-7 month time point, but not in the intervening period. In the cumulative cDC2 and cDC3 population, CD86 expression increased to a significantly higher level than controls at 6 weeks and 6-7 months (**Figure 7D**). Only cumulative cDC2 and cDC3 showed any significant changes to HLA-DR expression, with a significant increase over time, until the expression from 6 weeks onwards showed no significant difference to healthy controls (**Figure 7E**). Taken together these data show that COVID-19 elicited lasting alterations in PD-L1 and CD86 expression levels on DC subsets.

**Figure 7.**
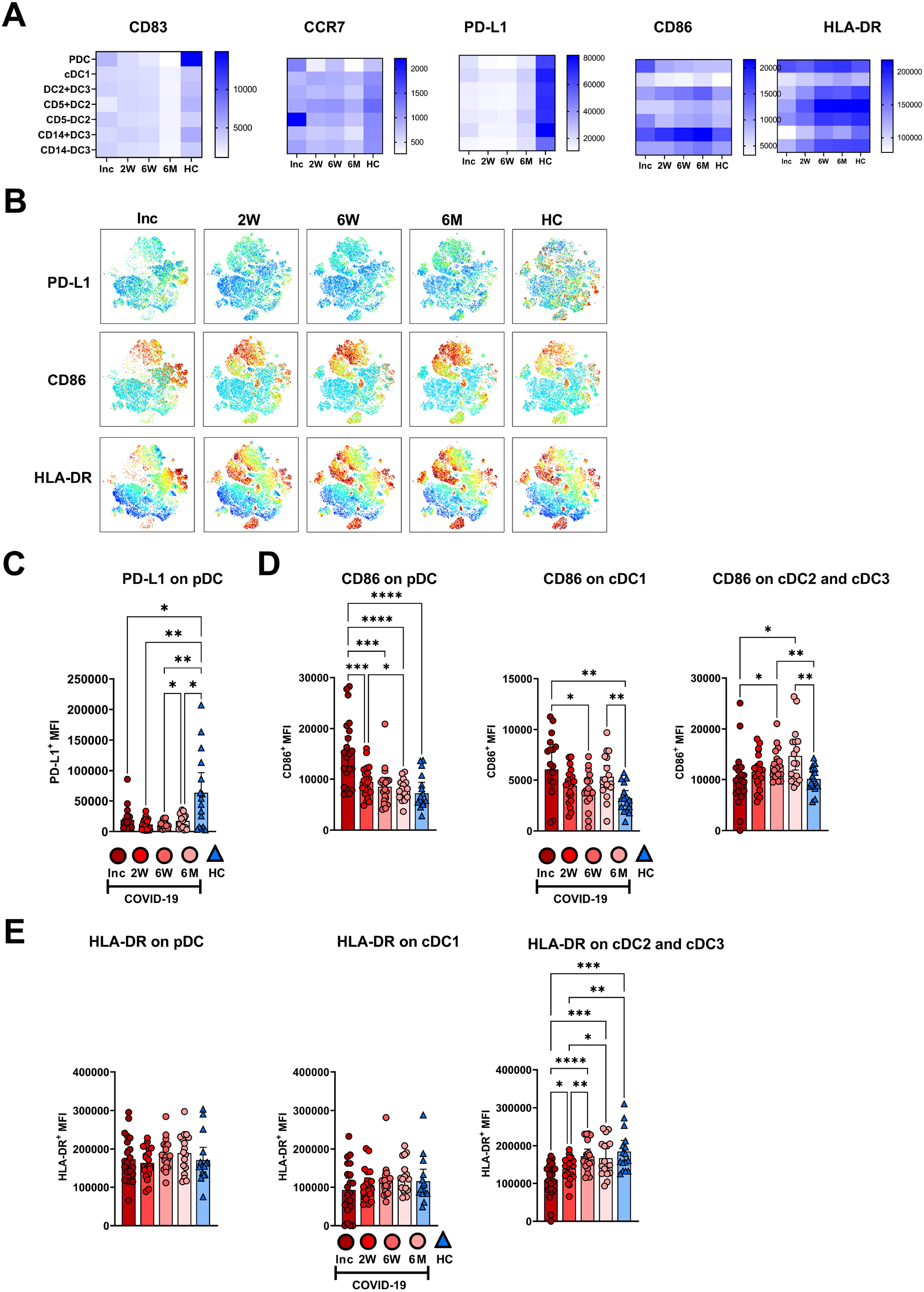
Altered phenotype of circulating dendritic cell subsets in COVID-19 patients. PBMCs collected from hospitalized COVID-19 patients (N = 21) and healthy controls (N = 16) over a 6-7 months period, were assessed for phenotypical changes. (**A**) Heatmaps showing the mean fluorescence intensity (MFI) of CD83, CCR7, PD-L1, CD86, and HLA-DR on different DC subsets. (**B**) tSNE plots showing CD88^-^ myeloid cells, colored according to intensity of PD-L1, CD86, and HLA-DR expression. (**C**) Representative graph of PD-L1 on pDC. MFI of (**D**) CD86 and (**E**) HLA-DR on DC subsets. Data is represented as mean with 95% Cl, with significance of *p ≤ 0.05, **p ≤ 0.01, ***p ≤ 0.001, ****p ≤ 0.0001., determined using Brown-Forsythe and Welch ANOVA tests. Inc = Study inclusion, 2W = 2 weeks, 6W = 6 weeks, 6M = 6-7 months, HC = healthy control.

### CRP at inclusion correlated with alterations in monocyte and DC subsets in COVID-19 patients

To determine if the initial inflammatory response to the SARS-CoV-2 infection affected the myeloid cell compartment in the COVID-19 patients, we assessed correlations between clinical parameters and myeloid cell populations. We found that the proportion of iMo at inclusion positively correlated with CRP (**Figure 8A**), and negative correlations were found with CRP and all DC subtypes, both at 2 and 6 weeks post-inclusion (**Figure 8B**).

**Figure 8.**
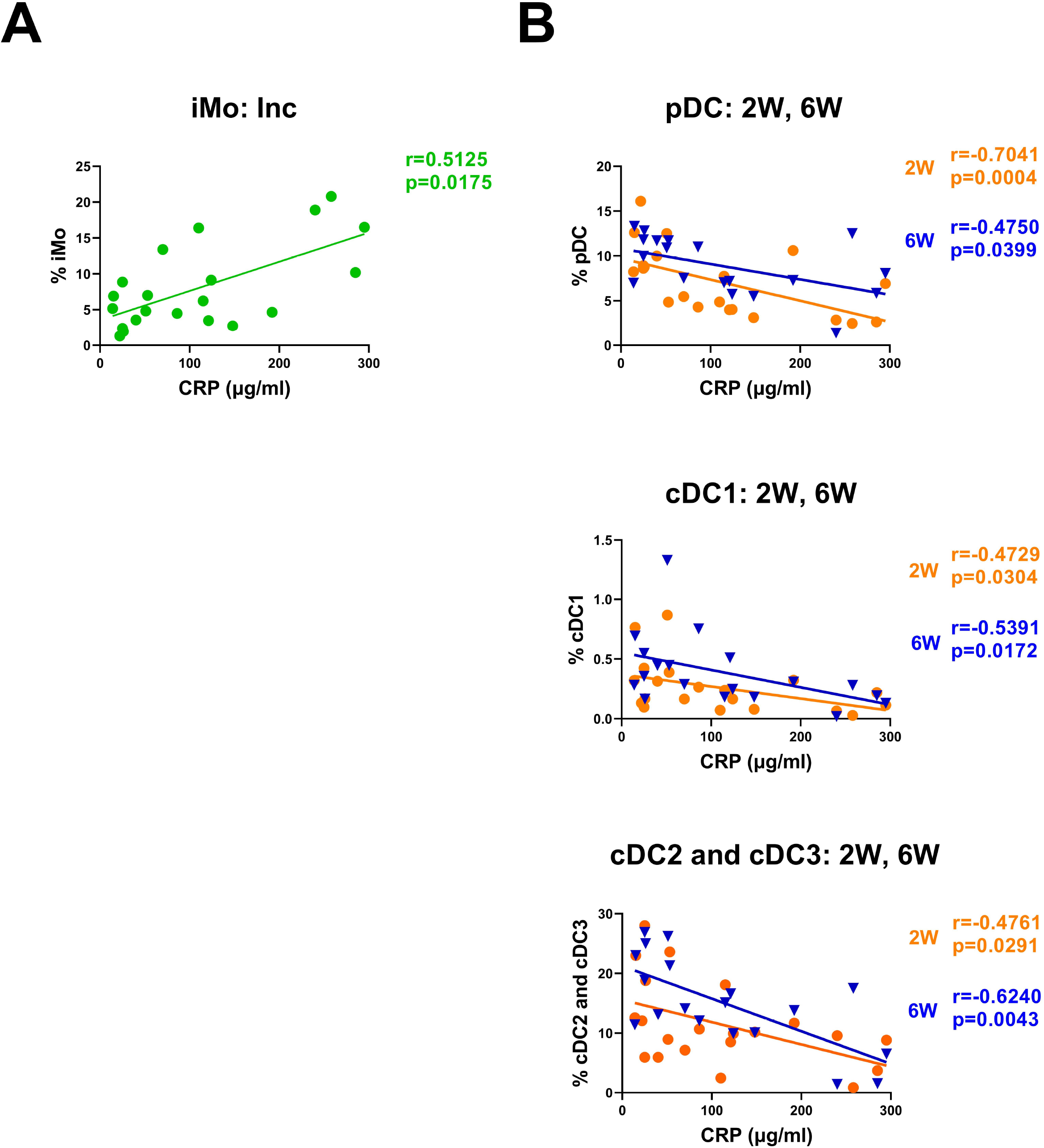
CRP levels correlated negatively with various monocyte and DC subsets. Bivariate analysis with Spearman’s correlation coefficient was performed using an array of clinical parameters and myeloid cell types. Significant correlations shown for (**A**) iMo at inclusion and (**B**) circulating DC subsets at 2 and 6 weeks post-inclusion, all against CRP. The p value and Spearman’s correlation coefficient (R) are shown for each analysis. N = 21. Inc = Study inclusion, 2W = 2 weeks, 6W = 6 weeks, 6M = 6-7 months, HC = healthy control.

## DISCUSSION

To address the lack of knowledge concerning myeloid cells in COVID-19 we have investigated the effects that SARS-CoV-2 infection exerts, both initially and long-term, on monocyte and DC subsets. Decreased frequencies of all DC subsets were found during acute COVID-19 with the levels subsequently returning to normal for all except the cDC2 and cDC3 combined subset, which still had higher levels at 6-7 months follow up. There were major alterations in monocyte subsets, with elevated levels of cMo that were still present at the long-term follow up. Notably, MDSC levels were still elevated at the 6-7 month timepoint. Additionally, our phenotypic assessment of functional markers showed elevated levels of CD86 on the monocyte subsets and MDSC at the final time point. On all cells HLA-DR expression was either unaffected or showed an initial decrease followed by recovery, except for iMo that had elevated HLA-DR from the second week and throughout the study period. Regarding CD86, the level at acute infection was elevated, especially on pDC, and high levels were sustained long term on cDC1 and cDC2/cDC3. HLA-DR expression was unaffected or low at inclusion on DC subsets and stayed low on the cDC2 and cDC3 combined subset. A lower expression of HLA-DR could be indicative of an initial suppressed functionality of amongst these DC.

Here we present a well-characterized, representative cohort, with clinical features commonly observed in hospitalized COVID-19 patients, as highlighted by marked lymphopenia, elevated serum CRP and LDH, and changes in the polymorphonuclear leukocytes, all in accordance with current knowledge ^32-35, 40^. Previous studies have illustrated a reduction of many of the myeloid cell subsets in COVID-19 ^23, 24, 27, 41^. Monocyte frequencies have been found to be lower during acute COVID-19 compared to patients in convalescence ^42^. In line with this, the levels of overall monocytes in the patients in our study were significantly decreased at inclusion, when compared to the later time points. The three subpopulations of monocytes are distinguished primarily by their expression of CD14 and CD16, but also by other cell surface markers, including CD36 and CCR2 for cMo, CCR5 and CD74 for iMo, and CXCR4 and CX3CR1 for nCMo. These markers aid in the wide range of functions of monocytes, including their antiviral activity ^43^. Our observed sustained increase in cMo in COVID-19 patients in comparison to healthy controls is consistent with previously published data ^23^. Key roles of cMo include tissue repair and anti-apoptotic functions ^44, 45^, and the elevated levels of these cells at 6-7 months could be part of the host resolving long-term effects of the SARS-CoV-2 infection. We saw an initial drastic drop in the level of ncMo, which recovered in convalescence, consistent with other reports ^23, 46^. Different patterns are seen in severe compared to mild COVID-19 in the levels of circulating iMo, with decreased levels in severe disease when measured during active disease ^24^. We found decreased levels of these cells compared to healthy controls and that they failed to reach the levels of the healthy controls before the 6-7 month follow up, indicating a long-lasting effect. In some COVID-19 patients with post-acute sequelae, i.e., long COVID, levels of ncMo and iMo were increased even 15 months post SARS-CoV-2 infection ^41^. From acute infection to convalescence, there was a decrease in cMo and an increase in ncMo ^47^. The decreased ncMo in severe COVID-19 patients during acute disease has been suggested to occur due to the recruitment of CD16^+^ myeloid cells to inflamed lung tissue during infection ^24, 48, 49^. In mouse models of acute lung injury and in acute respiratory distress syndrome, as seen in severe COVID-19, circulating monocytes have been shown to play a pivotal role in driving inflammation ^50^. Another factor affecting the levels of monocyte subsets could be virus-mediated cell death during COVID-19, since they all express the SARS-CoV-2 receptor, ACE2 ^42^, and there is evidence of viral antigens present within ncMo for over a year after initial infection ^41^. In addition, CD16^+^ monocytes can also be infected in an ACE2-independent manner, leading to inflammatory cell death ^51^.

Recent studies, through single-cell resolution methods, point to the expansion of suppressive myeloid cells in the blood as a hallmark of severe COVID-19 ^14, 23^, as confirmed by our data. These cells (MDSC) are a heterogeneous population of myeloid cells spanning granulocytes, macrophages and DC and these cells exist during normal conditions at diverse frequencies at different tissue ^52^. The levels of MDSC have been found to correlate to impaired T cell responses ^53, 54^. This sustained elevation suggests a long-lasting suppressive state, which we have found to be reflected in the suppressive T cell phenotypes defined in our COVID-19 cohort ^55^.

In line with other studies, we found that COVID-19 had a major impact on circulating DC subpopulations ^23^, with major reductions during active disease for pDC ^14^, cDC1 ^24^, and combined cDC2/cDC3 ^56^. Of note, the low level of pDC was independent of the study participants’ age and sex, which is in line with previous findings ^57^. Indeed, the reason for the reduction seen in pDC and cDC1 at inclusion could be due to several factors. A redirection of circulating pDC and cDC1 to lymphoid tissue has been well documented ^25, 58^ and cDC2 subtypes have been shown to be recruited into the lung tissue, though this is not seen for pDC and cDC1 ^24, 59^. The myeloid DC populations, cDC1 and cDC2/cDC3 returned to normal levels within 2 weeks. Of note, there were long-lasting effects, with higher levels of combined cDC2/cDC3 compared to healthy controls. A lasting effect with higher levels of HLA-DR^+^CD11c^+^ DC has been noted in individuals needing hospitalization, whereas normal levels of CD141^+^ DC, i.e., cDC1 were observed ^30^. Regarding pDC, we found these cells to be affected for a longer time, and to not fully return to the level found in healthy controls until the 6 week time point. Conversely, at 7 months in the study by Pérez-Gómez ^30^ the pDC levels were still not restored to normal.

Increased levels of CD163^+^CD14^+^ DC3 could play a role in COVID-19 mediated inflammation ^56^. In addition, the elevated levels of these cells could indicate an ongoing long-term systemic inflammation due to the damage inflicted by COVID-19 on the host. In addition to relocation from circulation, the observed decreases in DC might depend on the killing of immune cells by direct or indirect viral effects. Contrary to monocytes, circulating DC lack expression of ACE2. However, other DC receptors such as CD147 and DC-SIGN can facilitate SARS-CoV-2 binding and entry ^60, 61^ and this might enable their infection. This interaction has been shown to induce pro-apoptotic p53 pathways in pDC ^25^. Almost all DC functions are affected during COVID-19 ^25^ with probable consequences for the progression and outcome of the illness. Initial type I and III IFN responses are essential to the resolution of viral infection and are found to be impaired in severe COVID-19 ^62^. One possible explanation for this may be the reduction of pDC, which are a major source of type I IFN. Given the importance of DC for the initial activation of T cell responses, their depletion or tissue relocation could hamper the subsequent T cell response ^63^. Additionally, the long-lasting alteration in the DC compartment could explain the sustained T cell dysfunction seen following COVID-19 ^55, 64^.

In different diseases such as cancer and severe infections, the phenotype and functionality of circulating monocyte and DC subsets are altered ^65, 66^. Our phenotypic assessment for functional markers affecting cellular activation and suppression showed little to no decrease in CD86 expression during acute disease, on monocyte subsets and MDSC. This was followed by elevated CD86 levels on all these cells at the long-term follow up. Previous studies have shown that monocyte subsets, and especially iMo, have decreased CD86 during acute infection ^67, 68^. HLA-DR expression on monocyte subsets had a similar initial decrease as found for CD86, with a recovery to levels in healthy controls. The exception to this was iMo, that had elevated HLA-DR from the second week and throughout the study period. The decreased HLA-DR levels in acute infection for cMo, ncMo are in accordance with previous studies ^14, 46, 67, 69, 70^. The iMo have been found to have downregulated HLA-DR during severe acute COVID-19 ^14, 46, 69^, which differs to our observation of no effect on HLA-DR compared to healthy control at inclusion. For HLA-DR a more pronounced decrease has been connected to severe disease ^69^.

In severe COVID-19, HLA-DR has previously been found to be reduced across all circulating DC besides cDC1 ^23^. In our study HLA-DR was only significantly altered on cDC2/3 where it was reduced at early time points before returning to levels found in controls, which is in agreement with Marongiu et al. who showed a reduction of HLA-DR in cDC2 and cDC3 subsets ^71^. The observed initial low expression of HLA-DR on cDC1, cDC2/cDC3, cMo, and ncMo could be a sign of immunosuppression in these cells) ^26, 69, 72, 73^. Interestingly, the elevated levels on iMo found after a few weeks and throughout the study period could be an indication of an ongoing inflammatory environment due to residual effects of the disease ^8, 9^.

Regarding the costimulatory molecule CD86 on DC, we found an increased level of expression on pDC and cDC1 at inclusion but not cDC2/cDC3. The elevated CD86 levels on pDC have been documented during acute SARS-CoV-2 infection ^57^. However, other studies found no effect on pDC, nor did they find decreased CD86 expression on HLA-DR^+^CD11c^+^ DC ^27, 70^, or on other myeloid DCs subsets in acute disease ^7, 56^. The expression of CD86 returned to levels comparable to healthy controls quickly for pDC but remained elevated on cDC1 and cDC2 and cDC3 subsets at 6-7 months. This contrasts with a previously published study which showed long-term reduction in CD86 expression on pDC and HLA-DR^+^CD11c^+^ DC subsets in hospitalized COVID-19 patients ^30^. Elevated CD86 expression was found on ki67^+^ cDC2 and cDC3 subsets, i.e., a similar pattern as we have seen for these DC subsets, highlighting that the level seems to be connected to the time that they have been in circulation ^56^. CD86 is found to be elevated in cancer and chronic infections such as HIV-1, and the increased CD86 we see could be another indicator of ongoing, low-level systemic inflammation from lung repair following COVID-19 ^74^. The differences between our and other results could be due to the method of defining DC subsets, with many studies assessing CD14^-^HLA-DR^+^CD11c^+^ DC and not the different cDC subsets we have used in our study.

In our study we observed a consistently lower PD-L1 surface expression across pDC and cDC subsets, in keeping with gene and protein expression levels in hospitalized patients during acute disease ^30, 75, 76^. This decrease of PD-L1 has not been seen in other studies, rather an increased surface expression of PD-L1 has been seen when examining the bulk monocytes ^70, 77^, HLA-DR^+^CD11c^+^ DC, and pDC ^57, 70^. Interestingly, on cDC1 and cDC2 recently entering circulation (ki67^+^ cells), the levels of PD-L1 were decreased in COVID-19 patients compared to controls ^56^. In our data this reduced expression of PD-L1 persisted for the entire duration of the study, as was seen also in a previous study at 7 months post-COVID-19 ^30^. Furthermore, PD-L1 expression was reduced on monocyte subsets when compared to the healthy controls throughout the 6-7 month period of our study. The loss of PD-L1 might be due to shedding of soluble PD-L1, which is found to be elevated in the serum of COVID-19 patients ^75^. Of note, the severity of disease seems to correlate to the level of soluble PD-L1 in circulation ^75^. The lasting reduction that we observe in PD-L1 expression across all myeloid cell subsets requires further study to explore if it plays any role in COVID-19 pathogenesis.

Concerning the migratory receptor CCR7, there was little change in the DC and monocyte subsets during acute disease compared to healthy controls, while long-term effects included decreased CCR7 on cMo, pDC and the two cDC3 populations. This decrease is in accordance with previous findings for cDC at 7 months ^30^, but not for pDC, which had long-term increased CCR7 ^30^, though this study measured the percentage of positive cells whereas we measured level of expression. CD83 expression has been explored at gene level in DC in COVID-19 ^14, 78^, however, not much is known regarding surface protein expression. We did not find any major alterations in the surface expression of CD83 across DC, which concurs with Venet et al. ^76^. Overall, we identified marked alterations to the expression of surface markers across myeloid cell types following COVID-19. The long-term changes to the surface expression of these proteins on monocytes and DC may be due to epigenetic changes resulting from severe COVID-19 ^79^, possibly altering progenitors in the bone marrow.

COVID-19 severity has been linked to an array of clinical parameters such as the level of soluble urokinase plasminogen activator receptor (suPAR), CRP, and viral load ^80-83^. In this study, the viral load correlated positively with the CRP levels and negatively with anti-spike IgG and neutralizing SARS-CoV-2 antibody levels, at inclusion. In addition, the COVID-19 patients with high CRP levels had lower levels of anti-spike IgG and neutralizing antibodies at inclusion. This negative association between CRP and antibody levels early on during SARS-CoV-2 infection has, to our knowledge, not yet been made. It has previously been shown that higher levels of specific antibodies during convalescence correlated with initial higher CRP levels ^84^, though we did not see this. During acute disease, the levels of pDC, cDC2, and CD163^+^CD14^-^ cDC3 were negatively correlated to CRP levels ^56, 69^, which we confirm in our study to be the case for all circulating DC subsets, i.e., a faster DC recovery could be predicted by lower initial levels of CRP. Conversely, during mild disease a positive correlation of pDC proportions with CRP has been shown ^30^. The levels of all monocyte subsets were previously found to negatively correlate to CRP in acute disease ^85^, while we found a positive correlation for iMo levels in our study. The failure to induce a strong type I IFN response in COVID-19 leads to a prolonged high viral load and induces a highly inflammatory environment that includes raised CRP ^86^. This in turn might have long-term effects on monocyte and DC subsets, which our data strongly supports.

A potential limitation of our study is the imperfect age and sex matching of the controls to the patients. This is of particular importance for some DC subsets such as pDC, which are decreased in older (>40 years) healthy individuals, whereas there are no significant effects on cDC ^87, 88^. Given that the levels of pDC returned to the levels of healthy controls we do not believe this is an issue. While biological sex does not seem to play a role for the levels of MDSC in general, their levels could be influenced during disease. A study of MDSC in mild to severe COVID-19 found higher levels of monocytic MDSC in males than in females ^89^. The increase we found of MDSC in blood from hospitalized COVID-19 patients did not correlate with biological sex or age, however, our cohort was relatively small and did not contain cases of mild disease, so a larger cohort may be needed to observe these associations. The MDSC levels increase with age and highly elevated levels are seen in severe infections and cancers ^90-96^. For instance, there are increased conventional CD33^+^HLA-DR^−^CD45^+^ MDSC in older adults compared to younger adults. A general problem, not specific for our study, when comparing data from different COVID-19 studies is the definition of disease severity, that diverges depending on country. We defined severity according to the NIH guidelines, which are based largely on supplemental oxygen requirements ^31^, as opposed to the WHO scale ^97^.

In conclusion, given the long-lasting changes in the monocyte, DC and MDSC compartments, as seen in the altered frequencies of cell populations and expression levels of various surface markers, it is evident that COVID-19 imprints the development and fate of these cells. Further studies will be required to determine for exactly how long these alterations persist after severe COVID-19, and if they affect the type and quality of immune responses elicited against future infections.

## Authorship contribution

F.R.H., M.G., C.S., J.N., and H.W. conducted experiments. F.R.H., M.G., C.S., J.N., H.W., S.N., and M.L. analyzed the data. F.R.H., M.G., and M.L were involved in the writing of the initial manuscript, and F.R.H., M.G., C.S., J.N., A.N., A.J.H., M.H., J.S., S.N., and M.L. helped in the revision of the manuscript. M.L. designed the experiments. A.N., A.J.H., M.H., J.S., S.N., and M.L. were involved in establishing the cohort study. All authors read and approved the final manuscript.

## Acknowledgements

We are grateful for all patients and healthy donors who participated in this cohort study. We thank all individuals involved in the study, including coordinators, and all healthcare personnel from the Clinic of Infectious Diseases, and the Intensive Care Unit at the Vrinnevi Hospital, Norrköping, Sweden. Our sincere thanks to Annette Gustafsson for helping with the study coordination and sample collection. We would like to thank Mario Alberto Cano Fiestas and Robin Göransson for their assistance with sample processing. We appreciate the assistance from Jörgen Adolfsson and the Core Facility Flow Cytometry Unit at Linkoping University. Finally, we thank the staff at the Flow Cytometry Unit at Clinical Immunology and Transfusion Medicine, Region Östergötland.

## References

1. WHO. COVID-19 advice for the public: Getting vaccinated: World Health Organization; 2022 [Available from: https://www.who.int/emergencies/diseases/novel-coronavirus-2019/covid-19-vaccines/advice.

2. WHO. WHO Coronavirus (COVID-19) Dashboard Geneva2022 [updated 1 March. Website]. Available from: https://covid19.who.int/.

3. Moss P. The T cell immune response against SARS-CoV-2. Nat Immunol. 2022;23(2):186–93.

4. Chen Z, John Wherry E.w T cell responses in patients with COVID-19. Nat Rev Immunol. 2020;20(9):529–36.

5. Bassler K, Schulte-Schrepping J, Warnat-Herresthal S, Aschenbrenner AC, Schultze JL. The Myeloid Cell Compartment—Cell by Cell. Annual Review of Immunology. 2019;37(1):269–93.

6. Kapellos TS, Bonaguro L, Gemund I, Reusch N, Saglam A, Hinkley ER, et al. Human Monocyte Subsets and Phenotypes in Major Chronic Inflammatory Diseases. Front Immunol. 2019;10:2035.

7. Anbazhagan K, Duroux-Richard I, Jorgensen C, Apparailly F. Transcriptomic network support distinct roles of classical and non-classical monocytes in human. Int Rev Immunol. 2014;33(6):470–89.

8. Lee J, Tam H, Adler L, Ilstad-Minnihan A, Macaubas C, Mellins ED. The MHC class II antigen presentation pathway in human monocytes differs by subset and is regulated by cytokines. PLoS One. 2017;12(8):e0183594.

9. Tolouei Semnani R, Moore V, Bennuru S, McDonald-Fleming R, Ganesan S, Cotton R, et al. Human monocyte subsets at homeostasis and their perturbation in numbers and function in filarial infection. Infect Immun. 2014;82(11):4438–46.

10. Veglia F, Sanseviero E, Gabrilovich DI. Myeloid-derived suppressor cells in the era of increasing myeloid cell diversity. Nature Reviews Immunology. 2021;21(8):485–98.

11. Coates BM, Staricha KL, Koch CM, Cheng Y, Shumaker DK, Budinger GRS, et al. Inflammatory Monocytes Drive Influenza A Virus-Mediated Lung Injury in Juvenile Mice. J Immunol. 2018;200(7):2391–404.

12. Leon J, Michelson DA, Olejnik J, Chowdhary K, Oh HS, Hume AJ, et al. A virus-specific monocyte inflammatory phenotype is induced by SARS-CoV-2 at the immune–epithelial interface. Proceedings of the National Academy of Sciences. 2022;119(1):e2116853118.

13. Haschka D, Petzer V, Burkert FR, Fritsche G, Wildner S, Bellmann-Weiler R, et al. Alterations of blood monocyte subset distribution and surface phenotype are linked to infection severity in COVID-19 inpatients. Eur J Immunol. 2022.

14. Schulte-Schrepping J, Reusch N, Paclik D, Bassler K, Schlickeiser S, Zhang B, et al. Severe COVID-19 Is Marked by a Dysregulated Myeloid Cell Compartment. Cell. 2020;182(6):1419–40 e23.

15. Vanderbeke L, Van Mol P, Van Herck Y, De Smet F, Humblet-Baron S, Martinod K, et al. Monocyte-driven atypical cytokine storm and aberrant neutrophil activation as key mediators of COVID-19 disease severity. Nat Commun. 2021;12(1):4117.

16. Banchereau J, Steinman RM. Dendritic cells and the control of immunity. Nature. 1998;392(6673):245–52.

17. Collin M, Bigley V. Human dendritic cell subsets: an update. Immunology. 2018;154(1):3–20.

18. Ziegler-Heitbrock L, Ancuta P, Crowe S, Dalod M, Grau V, Hart DN, et al. Nomenclature of monocytes and dendritic cells in blood. Blood. 2010;116(16):e74–80.

19. Bao M, Liu YJ. Regulation of TLR7/9 signaling in plasmacytoid dendritic cells. Protein Cell. 2013;4(1):40–52.

20. Jongbloed SL, Kassianos AJ, McDonald KJ, Clark GJ, Ju X, Angel CE, et al. Human CD141+ (BDCA-3)+ dendritic cells (DCs) represent a unique myeloid DC subset that cross-presents necrotic cell antigens. J Exp Med. 2010;207(6):1247–60.

21. Yin X, Yu H, Jin X, Li J, Guo H, Shi Q, et al. Human Blood CD1c+ Dendritic Cells Encompass CD5high and CD5low Subsets That Differ Significantly in Phenotype, Gene Expression, and Functions. J Immunol. 2017;198(4):1553–64.

22. Peng Q, Qiu X, Zhang Z, Zhang S, Zhang Y, Liang Y, et al. PD-L1 on dendritic cells attenuates T cell activation and regulates response to immune checkpoint blockade. Nature Communications. 2020;11(1):4835.

23. Kvedaraite E, Hertwig L, Sinha I, Ponzetta A, Hed Myrberg I, Lourda M, et al. Major alterations in the mononuclear phagocyte landscape associated with COVID-19 severity. Proc Natl Acad Sci U S A. 2021;118(6).

24. Sanchez-Cerrillo I, Landete P, Aldave B, Sanchez-Alonso S, Sanchez-Azofra A, Marcos-Jimenez A, et al. COVID-19 severity associates with pulmonary redistribution of CD1c+ DCs and inflammatory transitional and nonclassical monocytes. J Clin Invest. 2020;130(12):6290–300.

25. Saichi M, Ladjemi MZ, Korniotis S, Rousseau C, Ait Hamou Z, Massenet-Regad L, et al. Single-cell RNA sequencing of blood antigen-presenting cells in severe COVID-19 reveals multi-process defects in antiviral immunity. Nat Cell Biol. 2021;23(5):538–51.

26. Winheim E, Rinke L, Lutz K, Reischer A, Leutbecher A, Wolfram L, et al. Impaired function and delayed regeneration of dendritic cells in COVID-19. PLOS Pathogens. 2021;17(10):e1009742.

27. Zhou R, To KK, Wong YC, Liu L, Zhou B, Li X, et al. Acute SARS-CoV-2 Infection Impairs Dendritic Cell and T Cell Responses. Immunity. 2020;53(4):864–77 e5.

28. Rydyznski Moderbacher C, Ramirez SI, Dan JM, Grifoni A, Hastie KM, Weiskopf D, et al. Antigen-Specific Adaptive Immunity to SARS-CoV-2 in Acute COVID-19 and Associations with Age and Disease Severity. Cell. 2020;183(4):996-1012.e19.

29. Calistri P, Amato L, Puglia I, Cito F, Di Giuseppe A, Danzetta ML, et al. Infection sustained by lineage B.1.1.7 of SARS-CoV-2 is characterised by longer persistence and higher viral RNA loads in nasopharyngeal swabs. Int J Infect Dis. 2021;105:753–5.

30. Pérez-Gómez A, Vitallé J, Gasca-Capote C, Gutierrez-Valencia A, Trujillo-Rodriguez M, Serna-Gallego A, et al. Dendritic cell deficiencies persist seven months after SARS-CoV-2 infection. Cellular & Molecular Immunology. 2021;18(9):2128–39.

31. NIH. COVID-19 Treatment Guidelines Panel. Coronavirus Disease 2019 (COVID-19) Treatment Guidelines. National Institutes of Health. 2020 [Available from: https://www.covid19treatmentguidelines.nih.gov/.

32. Huang C, Wang Y, Li X, Ren L, Zhao J, Hu Y, et al. Clinical features of patients infected with 2019 novel coronavirus in Wuhan, China. Lancet. 2020;395(10223):497–506.

33. Laing AG, Lorenc A, del Molino del Barrio I, Das A, Fish M, Monin L, et al. A dynamic COVID-19 immune signature includes associations with poor prognosis. Nature Medicine. 2020;26(10):1623–35.

34. Reusch N, De Domenico E, Bonaguro L, Schulte-Schrepping J, Baßler K, Schultze JL, et al. Neutrophils in COVID-19. Frontiers in Immunology. 2021;12.

35. Xie G, Ding F, Han L, Yin D, Lu H, Zhang M. The role of peripheral blood eosinophil counts in COVID-19 patients. Allergy. 2021;76(2):471–82.

36. Lourda M, Dzidic M, Hertwig L, Bergsten H, Palma Medina LM, Sinha I, et al. High-dimensional profiling reveals phenotypic heterogeneity and disease-specific alterations of granulocytes in COVID-19. Proc Natl Acad Sci U S A. 2021;118(40).

37. Thomopoulos TP, Rosati M, Terpos E, Stellas D, Hu X, Karaliota S, et al. Kinetics of Nucleocapsid, Spike and Neutralizing Antibodies, and Viral Load in Patients with Severe COVID-19 Treated with Convalescent Plasma. Viruses. 2021;13(9).

38. Röltgen K, Powell AE, Wirz OF, Stevens BA, Hogan CA, Najeeb J, et al. Defining the features and duration of antibody responses to SARS-CoV-2 infection associated with disease severity and outcome. Sci Immunol. 2020;5(54):eabe0240.

39. Mair F, Liechti T. Comprehensive Phenotyping of Human Dendritic Cells and Monocytes. Cytometry Part A. 2021;99(3):231–42.

40. Lourda M, Dzidic M, Hertwig L, Bergsten H, Medina LMP, Sinha I, et al. High-dimensional profiling reveals phenotypic heterogeneity and disease-specific alterations of granulocytes in COVID-19. Proceedings of the National Academy of Sciences. 2021;118(40):e2109123118.

41. Patterson BK, Francisco EB, Yogendra R, Long E, Pise A, Rodrigues H, et al. Persistence of SARS CoV-2 S1 Protein in CD16+ Monocytes in Post-Acute Sequelae of COVID-19 (PASC) up to 15 Months Post-Infection. Front Immunol. 2021;12:746021.

42. Rutkowska-Zapala M, Suski M, Szatanek R, Lenart M, Weglarczyk K, Olszanecki R, et al. Human monocyte subsets exhibit divergent angiotensin I-converting activity. Clin Exp Immunol. 2015;181(1):126–32.

43. Kapellos TS, Bonaguro L, Gemünd I, Reusch N, Saglam A, Hinkley ER, et al. Human Monocyte Subsets and Phenotypes in Major Chronic Inflammatory Diseases. Frontiers in Immunology. 2019;10.

44. Wong KL, Tai JJ-Y, Wong W-C, Han H, Sem X, Yeap W-H, et al. Gene expression profiling reveals the defining features of the classical, intermediate, and nonclassical human monocyte subsets. Blood. 2011;118(5):e16–e31.

45. Wong KL, Yeap WH, Tai JJY, Ong SM, Dang TM, Wong SC. The three human monocyte subsets: implications for health and disease. Immunologic Research. 2012;53(1):41–57.

46. Gatti A, Radrizzani D, Vigano P, Mazzone A, Brando B. Decrease of Non-Classical and Intermediate Monocyte Subsets in Severe Acute SARS-CoV-2 Infection. Cytometry A. 2020;97(9):887–90.

47. Rajamanickam A, Kumar NP, Pandiarajan AN, Selvaraj N, Munisankar S, Renji RM, et al. Dynamic alterations in monocyte numbers, subset frequencies and activation markers in acute and convalescent COVID-19 individuals. Sci Rep. 2021;11(1):20254.

48. Carvelli J, Demaria O, Vély F, Batista L, Chouaki Benmansour N, Fares J, et al. Association of COVID-19 inflammation with activation of the C5a–C5aR1 axis. Nature. 2020;588(7836):146–50.

49. Gatti A, Radrizzani D, Viganò P, Mazzone A, Brando B. Decrease of Non-Classical and Intermediate Monocyte Subsets in Severe Acute SARS-CoV-2 Infection. Cytometry Part A. 2020;97(9):887–90.

50. Dhaliwal K, Scholefield E, Ferenbach D, Gibbons M, Duffin R, Dorward DA, et al. Monocytes Control Second-Phase Neutrophil Emigration in Established Lipopolysaccharide-induced Murine Lung Injury. American Journal of Respiratory and Critical Care Medicine. 2012;186(6):514–24.

51. Junqueira C, Crespo A, Ranjbar S, de Lacerda LB, Lewandrowski M, Ingber J, et al. FcgammaR-mediated SARS-CoV-2 infection of monocytes activates inflammation. Nature. 2022.

52. Youn JI, Nagaraj S, Collazo M, Gabrilovich DI. Subsets of myeloid-derived suppressor cells in tumor-bearing mice. J Immunol. 2008;181(8):5791–802.

53. Serafini P, Mgebroff S, Noonan K, Borrello I. Myeloid-derived suppressor cells promote cross-tolerance in B-cell lymphoma by expanding regulatory T cells. Cancer Res. 2008;68(13):5439–49.

54. Veglia F, Perego M, Gabrilovich D. Myeloid-derived suppressor cells coming of age. Nat Immunol. 2018;19(2):108–19.

55. Govender M, Hopkins FR, Göransson R, Svanberg C, Shankar EM, Hjorth M, et al. Long-term T cell perturbations and waning antibody levels in individuals needing hospitalization for COVID-19. bioRxiv. 2022:2022.03.17.484640.

56. Winheim E, Rinke L, Lutz K, Reischer A, Leutbecher A, Wolfram L, et al. Impaired function and delayed regeneration of dendritic cells in COVID-19. PLoS Pathog. 2021;17(10):e1009742.

57. Severa M, Diotti RA, Etna MP, Rizzo F, Fiore S, Ricci D, et al. Differential plasmacytoid dendritic cell phenotype and type I Interferon response in asymptomatic and severe COVID-19 infection. PLoS Pathog. 2021;17(9):e1009878.

58. Liu C, Martins AJ, Lau WW, Rachmaninoff N, Chen J, Imberti L, et al. Time-resolved systems immunology reveals a late juncture linked to fatal COVID-19. Cell. 2021;184(7):1836–57 e22.

59. Campana P, Parisi V, Leosco D, Bencivenga D, Della Ragione F, Borriello A. Dendritic Cells and SARS-CoV-2 Infection: Still an Unclarified Connection. Cells. 2020;9(9).

60. Wang K, Chen W, Zhang Z, Deng Y, Lian JQ, D. P, et al. CD147-spike protein is a novel route for SARS-CoV-2 infection to host cells. Signal Transduct Target Ther. 2020;5(1):283.

61. Thepaut M, Luczkowiak J, Vives C, Labiod N, Bally I, Lasala F, et al. DC/L-SIGN recognition of spike glycoprotein promotes SARS-CoV-2 trans-infection and can be inhibited by a glycomimetic antagonist. PLoS Pathog. 2021;17(5):e1009576.

62. Hadjadj J, Yatim N, Barnabei L, Corneau A, Boussier J, Smith N, et al. Impaired type I interferon activity and inflammatory responses in severe COVID-19 patients. Science. 2020;369(6504):718–24.

63. Sallusto F, Lanzavecchia A. The instructive role of dendritic cells on T-cell responses. Arthritis Res. 2002;4 Suppl 3:S127–32.

64. Yang J, Zhong M, Zhang E, Hong K, Yang Q, Zhou D, et al. Broad phenotypic alterations and potential dysfunction of lymphocytes in individuals clinically recovered from COVID-19. J Mol Cell Biol. 2021;13(3):197–209.

65. Kiss M, Caro AA, Raes G, Laoui D. Systemic Reprogramming of Monocytes in Cancer. Front Oncol. 2020;10:1399.

66. Miller E, Bhardwaj N. Dendritic cell dysregulation during HIV-1 infection. Immunol Rev. 2013;254(1):170–89.

67. Laing AG, Lorenc A, Del Molino Del Barrio I, Das A, Fish M, Monin L, et al. A dynamic COVID-19 immune signature includes associations with poor prognosis. Nat Med. 2020;26(10):1623–35.

68. Arunachalam PS, Wimmers F, Mok CKP, Perera R, Scott M, Hagan T, et al. Systems biological assessment of immunity to mild versus severe COVID-19 infection in humans. Science. 2020;369(6508):1210–20.

69. Peruzzi B, Bencini S, Capone M, Mazzoni A, Maggi L, Salvati L, et al. Quantitative and qualitative alterations of circulating myeloid cells and plasmacytoid DC in SARS-CoV-2 infection. Immunology. 2020;161(4):345–53.

70. Parackova Z, Zentsova I, Bloomfield M, Vrabcova P, Smetanova J, Klocperk A, et al. Disharmonic Inflammatory Signatures in COVID-19: Augmented Neutrophils’ but Impaired Monocytes’ and Dendritic Cells’ Responsiveness. Cells. 2020;9(10).

71. Marongiu L, Protti G, Facchini FA, Valache M, Mingozzi F, Ranzani V, et al. Maturation signatures of conventional dendritic cell subtypes in COVID-19 suggest direct viral sensing. Eur J Immunol. 2022;52(1):109–22.

72. Mudd PA, Crawford JC, Turner JS, Souquette A, Reynolds D, Bender D, et al. Distinct inflammatory profiles distinguish COVID-19 from influenza with limited contributions from cytokine storm. Science Advances. 2020;6(50):eabe3024.

73. Spinetti T, Hirzel C, Fux M, Walti LN, Schober P, Stueber F, et al. Reduced Monocytic Human Leukocyte Antigen-DR Expression Indicates Immunosuppression in Critically Ill COVID-19 Patients. Anesth Analg. 2020;131(4):993–9.

74. Polidoro RB, Hagan RS, de Santis Santiago R, Schmidt NW. Overview: Systemic Inflammatory Response Derived From Lung Injury Caused by SARS-CoV-2 Infection Explains Severe Outcomes in COVID-19. Front Immunol. 2020;11:1626.

75. Sabbatino F, Conti V, Franci G, Sellitto C, Manzo V, Pagliano P, et al. PD-L1 Dysregulation in COVID-19 Patients. Front Immunol. 2021;12:695242.

76. Venet M, Sa Ribeiro M, Décembre E, Bellomo A, Joshi G, Villard M, et al. Severe COVID-19 patients have impaired plasmacytoid dendritic cell-mediated control of SARS-CoV-2-infected cells. medRxiv. 2021:2021.09.01.21262969.

77. Rutkowska E, Kwiecien I, Klos K, Rzepecki P, Chcialowski A. Intermediate Monocytes with PD-L1 and CD62L Expression as a Possible Player in Active SARS-CoV-2 Infection. Viruses. 2022;14(4).

78. Cai G, Du M, Bosse Y, Albrecht H, Qin F, Luo X, et al. SARS-CoV-2 Impairs Dendritic Cells and Regulates DC-SIGN Gene Expression in Tissues. Int J Mol Sci. 2021;22(17).

79. Cheong J-G, Ravishankar A, Sharma S, Parkhurst CN, Nehar-Belaid D, Ma S, et al. Epigenetic Memory of COVID-19 in Innate Immune Cells and Their Progenitors. bioRxiv. 2022:2022.02.09.479588.

80. Sadeghi-Haddad-Zavareh M, Bayani M, Shokri M, Ebrahimpour S, Babazadeh A, Mehraeen R, et al. C-Reactive Protein as a Prognostic Indicator in COVID-19 Patients. Interdisciplinary Perspectives on Infectious Diseases. 2021;2021:5557582.

81. Malik P, Patel U, Mehta D, Patel N, Kelkar R, Akrmah M, et al. Biomarkers and outcomes of COVID-19 hospitalisations: systematic review and meta-analysis. BMJ Evidence-Based Medicine. 2021;26(3):107–8.

82. Enocsson H, Idoff C, Gustafsson A, Govender M, Hopkins F, Larsson M, et al. Soluble Urokinase Plasminogen Activator Receptor (suPAR) Independently Predicts Severity and Length of Hospitalisation in Patients With COVID-19. Front Med (Lausanne). 2021;8:791716.

83. Chen W, Xiao Q, Fang Z, Lv X, Yao M, Deng M. Correlation Analysis between the Viral Load and the Progression of COVID-19. Computational and Mathematical Methods in Medicine. 2021;2021:9926249.

84. Guo J, Li L, Wu Q, Li H, Li Y, Hou X, et al. Detection and predictors of anti-SARS-CoV-2 antibody levels in COVID-19 patients at 8 months after symptom onset. Future Virol. 2021;0(0).

85. Emsen A, Sumer S, Tulek B, Cizmecioglu H, Vatansev H, Goktepe MH, et al. Correlation of myeloid-derived suppressor cells with C-reactive protein, ferritin and lactate dehydrogenase levels in patients with severe COVID-19. Scand J Immunol. 2022;95(1):e13108.

86. Ramasamy S, Subbian S. Critical Determinants of Cytokine Storm and Type I Interferon Response in COVID-19 Pathogenesis. Clinical Microbiology Reviews. 2021;34(3):e00299–20.

87. Vora R, Bernardo D, Durant L, Reddi D, Hart AL, Fell JM, et al. Age-related alterations in blood and colonic dendritic cell properties. Oncotarget. 2016;7(11):11913–22.

88. Jing Y, Shaheen E, Drake RR, Chen N, Gravenstein S, Deng Y. Aging is associated with a numerical and functional decline in plasmacytoid dendritic cells, whereas myeloid dendritic cells are relatively unaltered in human peripheral blood. Hum Immunol. 2009;70(10):777–84.

89. Falck-Jones S, Vangeti S, Yu M, Falck-Jones R, Cagigi A, Badolati I, et al. Functional monocytic myeloid-derived suppressor cells increase in blood but not airways and predict COVID-19 severity. J Clin Invest. 2021;131(6).

90. Chi N, Tan Z, Ma K, Bao L, Yun Z. Increased circulating myeloid-derived suppressor cells correlate with cancer stages, interleukin-8 and -6 in prostate cancer. Int J Clin Exp Med. 2014;7(10):3181–92.

91. Huang A, Zhang B, Yan W, Wang B, Wei H, Zhang F, et al. Myeloid-derived suppressor cells regulate immune response in patients with chronic hepatitis B virus infection through PD-1-induced IL-10. J Immunol. 2014;193(11):5461–9.

92. Lv J, Zhao Y, Zong H, Ma G, Wei X, Zhao Y. Increased Levels of Circulating Monocytic- and Early-Stage Myeloid-Derived Suppressor Cells (MDSC) in Acute Myeloid Leukemia. Clin Lab. 2021;67(3).

93. Sacchi A, Grassi G, Notari S, Gili S, Bordoni V, Tartaglia E, et al. Expansion of Myeloid Derived Suppressor Cells Contributes to Platelet Activation by L-Arginine Deprivation during SARS-CoV-2 Infection. Cells. 2021;10(8).

94. Tcyganov EN, Hanabuchi S, Hashimoto A, Campbell D, Kar G, Slidel TW, et al. Distinct mechanisms govern populations of myeloid-derived suppressor cells in chronic viral infection and cancer. J Clin Invest. 2021;131(16).

95. Yaseen MM, Abuharfeil NM, Darmani H. Myeloid-derived suppressor cells and the pathogenesis of human immunodeficiency virus infection. Open Biol. 2021;11(11):210216.

96. Bowdish DM. Myeloid-derived suppressor cells, age and cancer. Oncoimmunology. 2013;2(7):e24754.

97. WHO Working Group on the Clinical Characterisation and Management of COVID-19 Infection-. A minimal common outcome measure set for COVID-19 clinical research. Lancet Infect Dis. 2020;20(8):e192–e7.

