## Supplementary File for "Lasting alterations in monocyte and dendritic cell subsets in individuals after hospitalization for COVID-19"

**Hopkins et al Supplementary file**

**Supplementary Table 1: Antibodies used for myeloid cell subset characterization**

| Marker | Fluorochrome | Company | Clone | Dilution |
|---|---|---|---|---|
| CD1c | PE-Cy7 | BioLegend | L161 | 1/80 |
| CD56 | Spark NIR 685 | BioLegend | 5.1H11 | 1/20 |
| CD163 | PE | BioLegend | GHI/61 | 1/20 |
| CD14 | BV510 | BD | MPhi9 | 1/40 |
| CD172a | BV650 | BD | SE5A5 | 1/640 |
| CD303 | BV786 | BD | V24-785 | 1/160 |
| HLA-DR | APC-H7 | BD | G46-6 | 1/40 |
| CADM1 | FITC | MBL | 3E1 | 1/320 |
| CD141 | BV605 | BD | 1A4 | 1/80 |
| FcεR1α | BB700 | BD | AER-37 | 1/160 |
| CD19 | AF647 | BioLegend | HIB19 | 1/80 |
| CD5 | BV711 | BD | UCHT2 | 1/80 |
| CD88 | AF700 | Bio-rad | P12/1 | 1/10 |
| CD11c | APC | BD | B-ly6 | 1/10 |
| CD3 | NovaBlue 610 | Phitonex* | UCHT1 | 1/40 |
| CD83 | BV421 | BioLegend | HB15e | 1/780 |
| CD86 | PE-Cy5 | BD | IT2.2 | 1/20 |
| CCR7 | BV750 | BioLegend | G043H7 | 1/20 |
| PD-L1 | BV480 | BD | MIH1 | 1/80 |
| CD16 | BV570 | BioLegend | 3G8 | 1/40 |
| Viability | LD Aqua | ThermoFisher | | 1/40 |

*now part of ThermoFisher

### Supplementary Figure 1

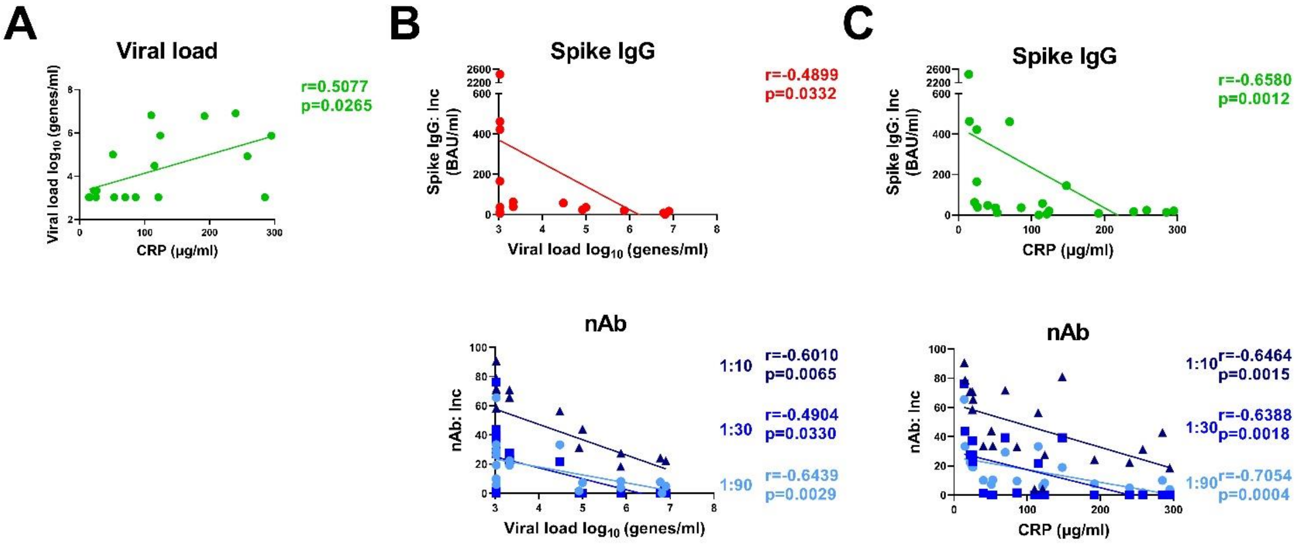


**Supplementary Figure 1**

**Correlations of clinical parameters in COVID-19 patients.** Samples were obtained from COVID-19 patients needing hospitalization (N=21). Bivariate analysis with Spearman's correlation coefficient was performed on clinical parameters at study inclusion. Significant correlations are shown for (**A**) viral load, and (**B**) spike IgG and neutralizing antibodies (nAb) against viral load, and (**C**) spike IgG and neutralizing antibodies against CRP. The p value and Spearman's correlation coefficient (R) shown for each analysis.

### Supplementary Figure 2

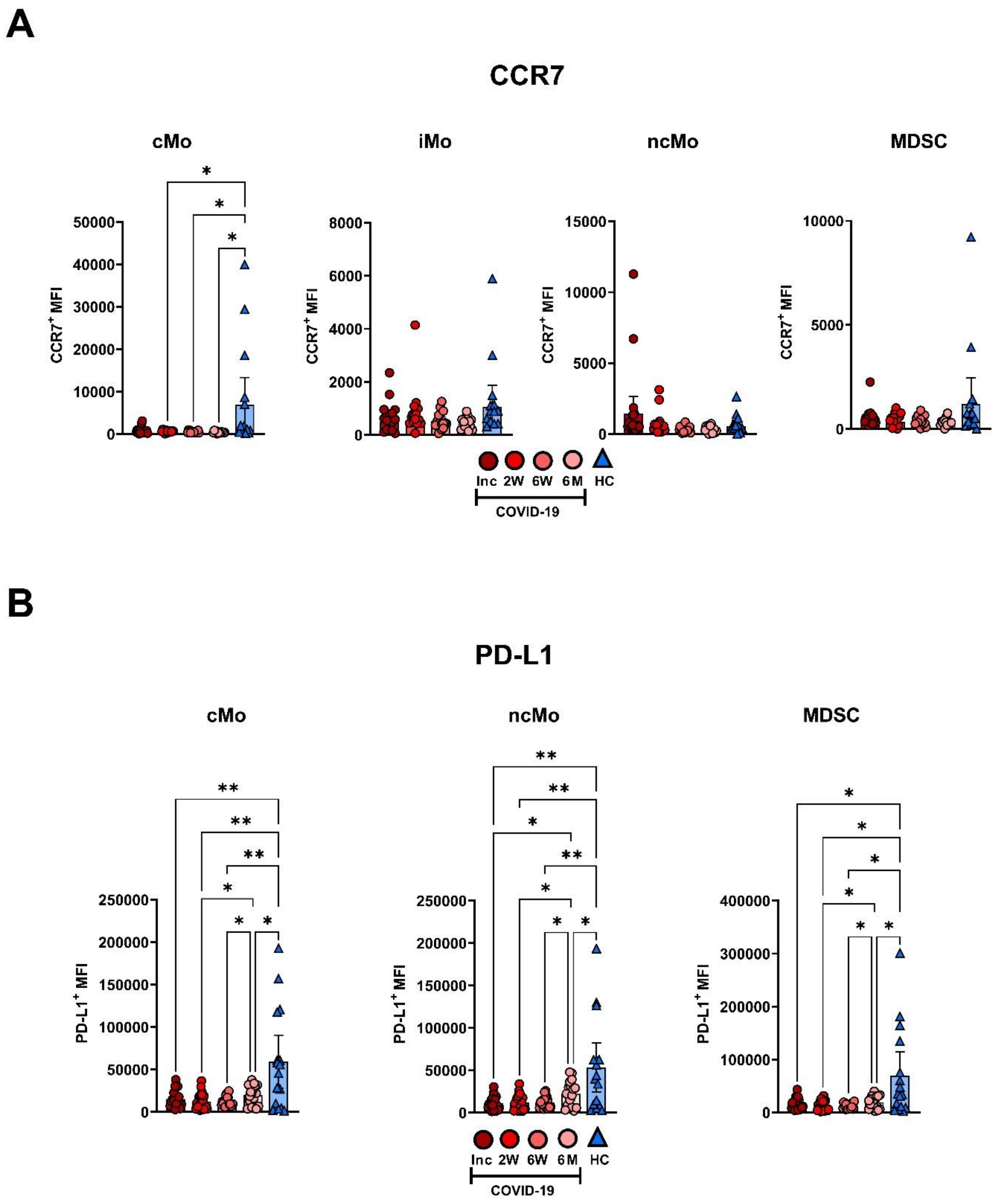


**Supplementary Figure 2: Altered phenotype in maturation and co-stimulation of blood monocyte cells subsets in COVID-19 patients.** PBMCs were obtained from COVID-19 patients needing hospitalization (N=21) and healthy controls (N=16) over a 6-7-month period. Data points represent Mean fluorescence intensity (MFI) of (**A**) CCR7 and (**B**) PD-L1 on monocytes and MDSC. Data is represented as mean with 95% Cl, with significance of *p≤0.05, **p≤0.01, ***p≤0.001, ****p≤0.0001., determined using Brown-Forsythe and Welch ANOVA tests. Inc = Inclusion in study at the hospital, 2W = 2 weeks, 6W = 6 weeks, 6M = 6-7 months, HC = healthy control.

### Supplementary Figure 3

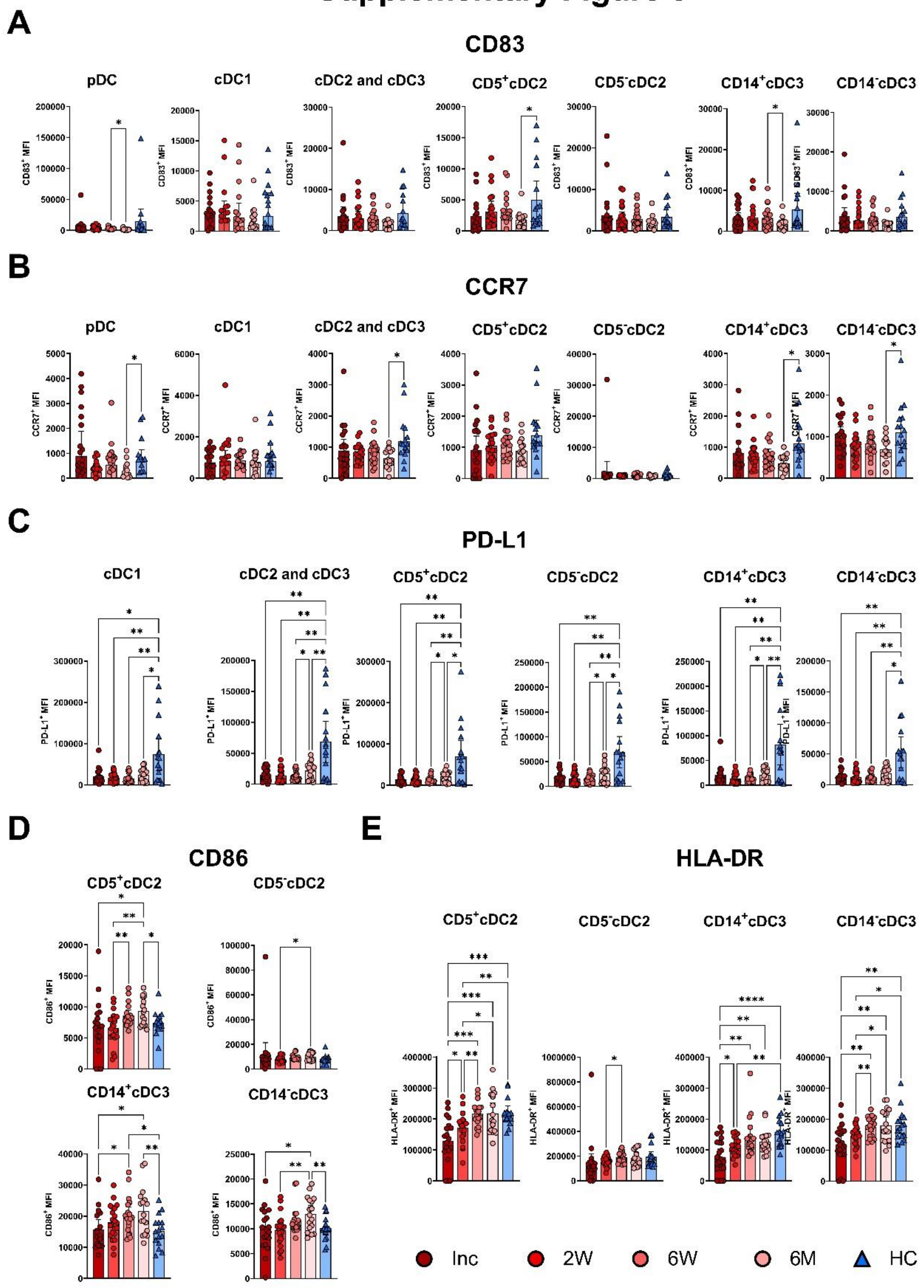


**Supplementary Figure 3: Changes in maturation and costimulatory markers on blood dendritic cells subsets in COVID-19 patients.** PBMCs from COVID-19 patients requiring hospitalization (N=21) and healthy controls (N=16) over a 6-7-month period were evaluated for phenotypical changes. Mean fluorescence intensity (MFI) of (**A**) CD83, (**B**) CCR7, (**C**) PD-L1, (**D**) CD86 and (**E**) HLA-DR on DC subsets. Data is represented as mean with 95% CI, with significance of *p≤0.05, **p≤0.01, ***p≤0.001, ****p≤0.0001., determined using Brown-Forsythe and Welch ANOVA tests. Inc = Inclusion in study at the hospital, 2W = 2 weeks, 6W = 6 weeks, 6M = 6-7 months, HC = healthy control.
